# SARS-CoV-2 mRNA vaccine is re-adenylated *in vivo*, enhancing antigen production and immune response

**DOI:** 10.1101/2022.12.01.518149

**Authors:** Paweł S Krawczyk, Olga Gewartowska, Michał Mazur, Wiktoria Orzeł, Katarzyna Matylla-Kulińska, Sebastian Jeleń, Paweł Turowski, Tomasz Śpiewla, Bartosz Tarkowski, Agnieszka Tudek, Aleksandra Brouze, Aleksandra Wesołowska, Dominika Nowis, Jakub Gołąb, Joanna Kowalska, Jacek Jemielity, Andrzej Dziembowski, Seweryn Mroczek

## Abstract

Though mRNA vaccines against COVID-19 have revolutionized vaccinology and have been administered in billions of doses, we know incredibly little about how mRNA vaccines are metabolized *in vivo*. Here we implemented enhanced nanopore Direct RNA sequencing (eDRS), to enable the analysis of single Moderna’s mRNA-1273 molecules, giving *in vivo* information about the sequence and poly(A) tails.

We show that mRNA-1273, with all uridines replaced by N1-methylpseudouridine (mΨ), is terminated by a long poly(A) tail (~100 nucleotides) followed by an mΨCmΨAG sequence. In model cell lines, mRNA-1273 is swiftly degraded in a process initiated by the removal of mΨCmΨAG, followed by CCR4-NOT-mediated deadenylation. In contrast, intramuscularly inoculated mRNA-1273 undergoes more complex modifications. Notably, mRNA-1273 molecules are re-adenylated after mΨCmΨAG removal. Detailed analysis of immune cells involved in antigen production revealed that in macrophages, after mΨCmΨAG removal, vaccine mRNA is very efficiently re-adenylated, and poly(A) tails can reach up to 200A. In contrast, in dendritic cells, vaccine mRNA undergoes slow deadenylation-dependent decay. We further demonstrate that enhancement of mRNA stability in macrophages is mediated by TENT5 poly(A) polymerases, whose expression is induced by the vaccine itself. Lack of TENT5-mediated re-adenylation results in lower antigen production and severely compromises specific immunoglobulin production following vaccination.

Together, our findings provide an unexpected principle for the high efficacy of mRNA vaccines and open new possibilities for their improvement. They also emphasize that, in addition to targeting a protein of interest, the design of mRNA therapeutics should be customized to its cellular destination.

## Main

The worldwide pandemic thrust vaccine development, and specifically development of mRNA-based vaccines. Thankfully for humankind, mRNA vaccines were deployed at an almost unimaginable speed and at worldwide scale.

Though pressed into immediate action, the mRNA vaccine field is still at very early stages. As this method of vaccine design was entirely new, it is essentially unknown how cellular and organismal systems respond to administration of an mRNA vaccine. Successful targeting of an mRNA vaccine, such as that against the SARS-CoV-2 spike protein, could be vastly improved by understanding the systems responsible for disseminating the vaccine in the body; uptake of the exogenous mRNA in different cell types; how or if mRNA inside the cell is maintained or metabolized; and how the protein product of the mRNA is used to build an immune reaction.

Clarification and harnessing of even some of these factors could make a significant difference in the efficacy and utility of mRNA vaccines. These factors could be of critical importance for reducing the ongoing burden of COVID-19, and also be broadly applied to improve the action and efficacy of many viral agents. The most potent vaccines for combating COVID-19 (BNT162b2 and mRNA-1273) to date are based on *in vitro*-transcribed (IVT) mRNA^1,2^. mRNA vaccines are administered encapsulated in lipid nanoparticles (LNPs) and enter cells mainly by endocytic pathways^3,4^. A fraction of these payloads is released from endosomes to the cytosol, where protein production occurs. Intramuscularly administered mRNA vaccines transfect neighboring resident immune cells, such as antigen-presenting dendritic cells (DCs) and macrophages. These cells become activated and migrate to neighboring lymph nodes^5–8^. LNPs, as well as secreted antigens produced in muscles, are also transported to local lymph nodes, where they are taken up by macrophages and resident DCs^8,9^. Although not directly demonstrated, it is likely that vaccine antigens transported with lymph and/or locally produced in the lymph nodes are stored on the surface of follicular DCs and induce a humoral immune response. DCs, which are central for adaptive immunity, endocytose antigens produced by other cells and translate vaccine mRNAs^9^. Peptide antigens are presented in association with class I and class II MHC molecules to T cells to induce effector CD8^+^ and CD4^+^ T cells^4,8^. The latter include helper (Th1) and follicular helper T (Tfh1) cells that are involved in the regulation of antiviral immunity in the periphery and development of humoral immune response in the follicles, respectively^8^. Many studies on mRNA vaccines have focused on DCs^10,11^. However, though macrophages/monocytes are the main cell population that take up mRNA vaccines^5,8^, their role in mRNA vaccine efficacy is not well defined.

Therapeutic mRNA resembles normal mRNA, having both a cap structure and poly(A) tail^3^, but it is generated through *in vitro* transcription, usually by T7 polymerase on a DNA template. A breakthrough in the development of RNA therapeutics came with the discovery that replacement of uridine with N1-methylpseudouridine (mΨ) decreases the innate immune response and enhances mRNA stability^12–15^. Although most currently synthesized mRNA therapeutics contain this modification, exceptions show it is not essential^16,17^. The two worldwide-approved mRNA vaccines, Bnt162b2^2,18^ and mRNA-1273^1,19^, encoding coronaviral spike protein (S protein), are produced using IVT and have all uridines replaced with mΨ, as well as a similar 5’ cap that is incorporated co-transcriptionally for Bnt162b2 and post-transcriptionally for mRNA-1273. However, they differ in their UTR sequences and 3’ tails^20^. Bnt162b2 vaccine has a composite poly(A) tail, with 30As followed by 10 other nucleotides, and then 70 additional As^18^, whereas mRNA-1273 has a poly(A) tail of undisclosed length. Both vaccines are formulated into nanoparticles with a similar lipid composition. The immune response was characterized for primary, secondary, and third dose vaccinations^19,21^, showing potent protection against the original variants, but also against variants of concern (VOC).

Knowledge about *in vivo* and *in-cell* metabolism of mRNA vaccines is extremely limited and is largely based on extrapolation from experiments in established cell lines with generic mRNAs. The paradigm extrapolated from data obtained on endogenous transcripts (or reporters) asserts that the stability of therapeutic mRNAs is determined by the rate of poly(A) tail removal (deadenylation). Notably, almost nothing is known regarding the metabolism of vaccines’ poly(A) tails. Herein, we implement nanopore direct RNA sequencing for a detailed dissection of mRNA-1273, revealing the complexity of its metabolism. Importantly, we show that mRNA-1273 is stabilized in macrophages by TENT5-mediated re-adenylation. We further demonstrate that TENT5A enhances the expression of the mRNA-1273 antigen *in vivo*, which is essential for an efficient immune response. Notably, in the past, the action of the endogenous polyadenylation machinery on mRNA therapeutics has not been reported or even considered.

### Direct RNA sequencing of mRNA vaccines

Studies of therapeutic mRNA metabolism face methodological challenges and, consequently, knowledge about the *in vivo* metabolism of mRNA vaccines is sparse. Approaches based on sequencing are usually indirect and rely on analysis of PCR-amplified cDNA. For poly(A) tails, such studies are error-prone, due to polymerase slipping on homopolymers. Similar problems concern the quality control of mRNA therapeutics, especially their poly(A) tails. Direct RNA sequencing (DRS, Oxford Nanopore Technologies^18^) has the potential to overcome technical barriers, as it is based on detecting electric current during the passage of single ssRNA molecules through a protein pore. Thus, DRS can provide reliable information on RNA composition with single-molecule resolution. However, the replacement of uridine with mΨ in mRNA therapeutics poses additional challenges, as this substitution perturbs current signal recorded during DRS and can result in imprecise translation into sequence (basecalling)^23^.

Analysis of intact mRNA-1273 using a standard DRS pipeline (**Fig. 1a**) showed that a significant proportion of aligned reads covered the full-length vaccine (**Fig. 1b**, blue), showing promise for comprehensive analysis of mRNA therapeutics via DRS. However, these experiments confirmed that mΨ indeed affects the basecalling process [Extended Data (ED) Fig 1a)]. Only 35.74% of reads aligned to the reference vaccine sequence, and those aligned reads had only 74.1% identity to the reference. Thus, the current data analysis pipeline is presently unsuitable for mΨ modified mRNAs.

**Figure 1.**
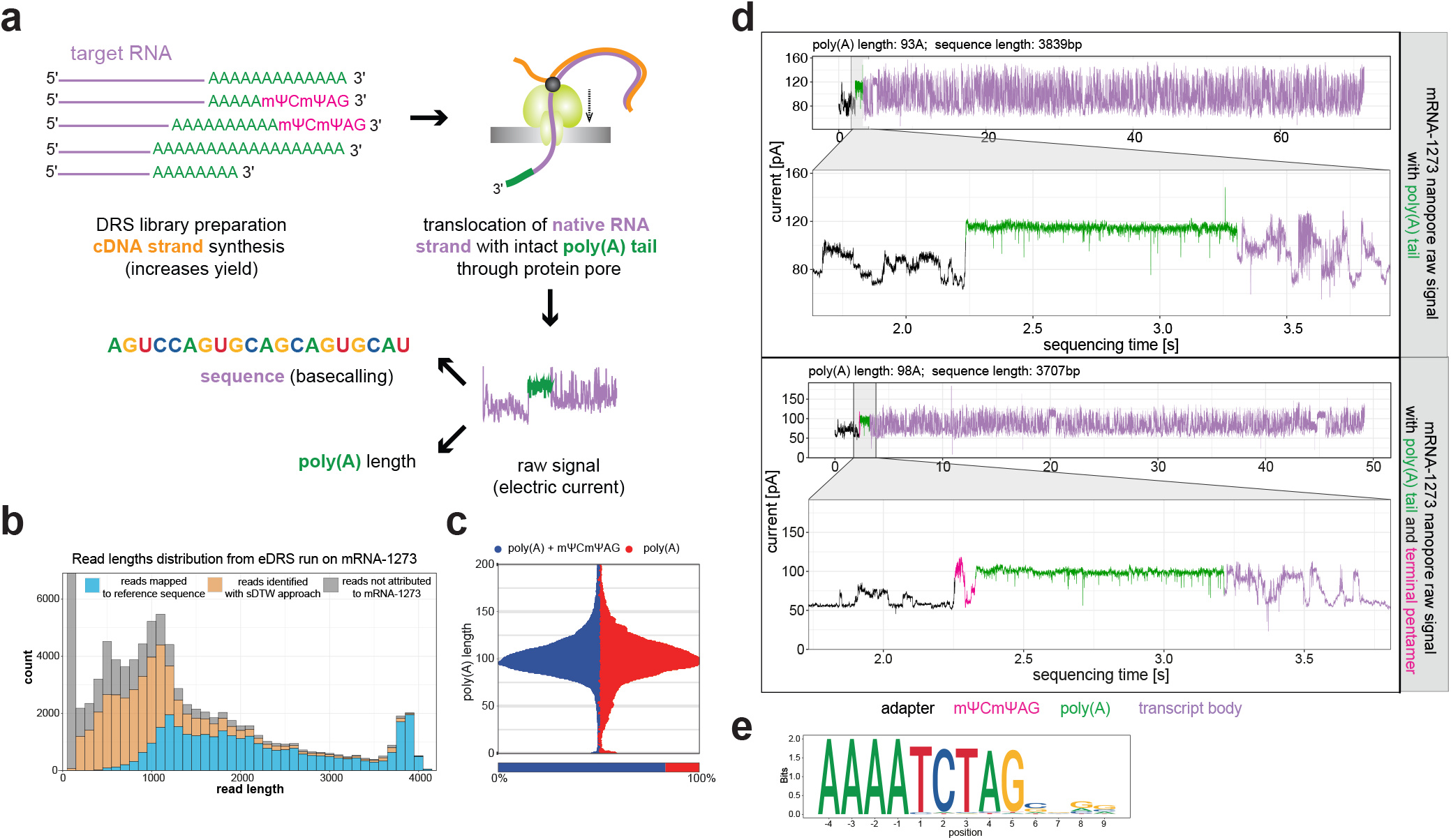
Enhanced Direct RNA Sequencing (eDRS) of mRNA-1273. **a**, Overview of DRS. Library with DNA adaptor ligated to native mRNAs is loaded on the flowcell with protein pores (yellow) embedded in the membrane. RNA molecules are translocated from the 3’ end through the pore due to electric potential and activity of the motor protein (grey dot). Passing through the pore causes perturbation in electric current readout, further translated (basecalled) into the sequence, and subsequently used for the poly(A) length calculation. **b**, Read lengths distribution of analyzed DRS data. Blue– reads mapped to mRNA-1273 after standard basecalling by Guppy. Yellow– reads identified as mRNA-1273 using Dynamic Time Warping (sDTW) approach. Grey – reads not attributed to mRNA-1273 (too low quality). Bin width = 100, stacked histogram. **c**, Distribution of poly(A) lengths for mRNA-1273 with (blue) and without (red) 3’ terminal mΨCmΨAG. Bottom panel – fraction of reads in each group is indicated. **d**, Representative raw signals from eDRS showing mRNA-1273 with and without 3’ terminal mΨCmΨAG, bottom and top panels, respectively. Signal from poly(A) is zoomed in for visualization purposes. Purple– transcript body, green – poly(A), black – adaptor, pink - mΨCmΨAG. **e**, Sequence logo of mRNA-1273 poly(A) 3’end obtained from 3’ RACE-seq. Last 4 adenosines from poly(A) tail are shown, followed by terminal nucleotides.

To improve our ability to identify desired reads, we developed a subsequence Dynamic Time Warping (sDTW) approach (ED Fig. 1b], to identify ionic current signatures specific to the 3’ end of mRNA-1273. This method, in principle, is similar to targeted nanopore sequencing^24^ and independent of basecalling and mapping. Compared to mapping alone, a combination of sDTW with mapping allowed the identification of twice as many sequences (ED Fig. 1c), validating the feasibility and promise of this approach. It was especially valuable for identifying shorter, lower-quality 3’-terminal reads (ED Fig. 1d), which would have been missed otherwise.

Next, we focused on improving accuracy in the determination of vaccine poly(A) tails, which are essential for mRNA stability and translation. Initially, we employed the nanopolish-polya algorithm to identify and measure poly(A) tails^25^, with good accuracy (ED Fig. 1e). This study revealed that mRNA-1273 has a poly(A) tract of ~100 adenosines (**Fig. 1c,** ED Fig. 1f). However, visual inspection of the raw current revealed unexpected perturbation between the poly(A) tail and the adaptor used for library preparation (**Fig. 1d**, bottom panel, green). Such a signal suggests the presence of terminal non-adenosine residue(s) in the mRNA-1273 vaccine. To verify whether this signature is specific to the 3’end of mRNA-1273, and not introduced during library preparation, we repeated the experiments with a vaccine RNA whose 3’ end was enzymatically extended with inosine nucleotides (I-tailing), followed by direct RNA sequencing with a custom protocol enabling the incorporation of such RNAs into the library. This analysis reproduced the same current perturbation (ED Fig. 1g).

Finally, Rapid Amplification of cDNA 3’ End (3’-RACE) Illumina sequencing revealed the presence of TCTAG (mΨCmΨAG in vaccine mRNA) (**Fig. 1e**), which likely represents the residue of restriction enzyme cleavage of the DNA template. For quantitative studies of mRNA-1273, we generated an enhanced DRS pipeline (eDRS) incorporating sDTW and a modified algorithm for the detection of poly(A) tails with mΨCmΨAG. Analysis of intact vaccine RNA showed that the major fraction of vaccine RNA molecules had an intact 3’ end. We observed that roughly 20% of the analyzed reads lacked the mΨCmΨAG sequence (**Fig. 1c**, red, bottom panel) and were characterized by a broader distribution of poly(A) lengths (**Fig. 1c**, red, upper panel). We propose that the eDRS pipeline is well suited to the analysis of mRNA vaccines and, in principle, could serve as a quality control step for mRNA therapeutics.

### Rapid deadenylation of mRNA-1273 in model cell lines

We then used eDRS to monitor stability and poly(A) tail status in cells typically used for pre-clinical analysis. We first determined the optimal amount of vaccine RNA: non-toxic to cells, allows efficient protein production, and detectable in bulk RNA with eDRS (**Fig. 2a**, ED Fig. 2ab). Then, HEK293T and A549 cells were transfected and cultured for up to 72 h, followed by RNA extraction and DRS. eDRS efficiently separated vaccine mRNA from endogenous mRNAs (ED Fig. 2c), allowing recovery of up to twice as many reads, compared to the standard basecalling/mapping (ED Fig. 2d). Analysis of poly(A) tails revealed that mΨCmΨAG was swiftly removed in both cell lines, followed by deadenylation (**Fig. 2bc,** ED Fig. 2e); this was observed as a reduction in mean poly(A) length with time, accompanied by a decreased number of vaccines reads. Concordantly with the RNA decay, spike protein levels also decreased with time (**Fig. 2d**), confirming direct relation between RNA stability and protein production.

**Figure 2.**
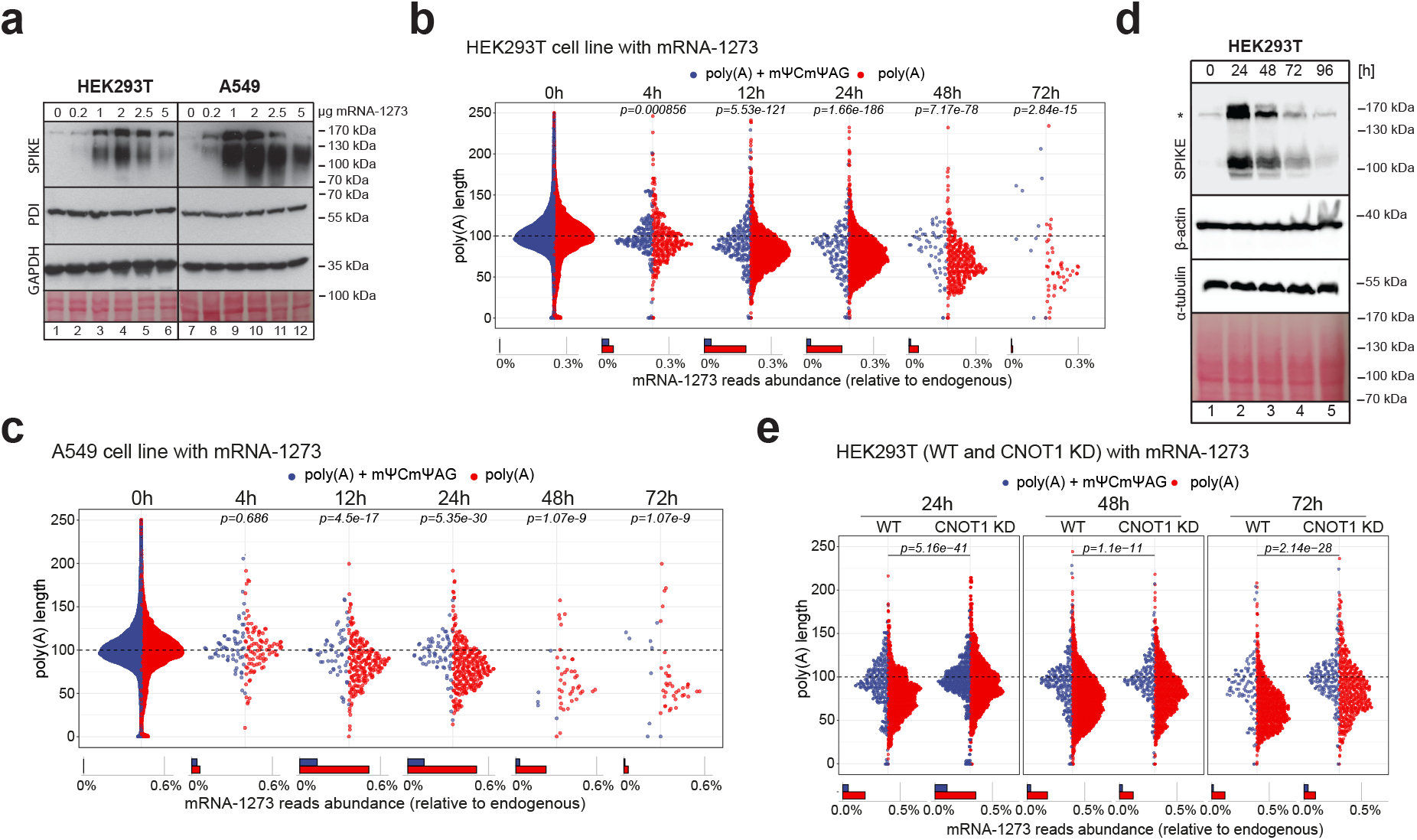
Stability of mRNA-1273 in model cell lines is determined by the deadenylation rate. **a,** Western blots showing efficient translation of mRNA-1273 in HEK293T and A549 cells with varying amounts of mRNA-1273 LNP. Asterisk indicate unspecific band. **b-c**, mRNA-1273 poly(A) length distribution in HEK293T (**b)**and A549 (**c)**cells treated with the vaccine for up to 72 h. Reads are divided into ones with (blue) and without (red) mΨCmΨAG sequence. Lower panel shows the relative abundance of each group. Average distributions for 1 (A549) or 2 (HEK293T) replicates are shown. P.values were calculated using Wilcoxon test, p. value adjustment with the Benjamini-Hochberg method. **d,**Expression of mRNA-1273 in HEK293T cells up to 96 h after mRNA-1273 delivery. Western Blot on spike protein. Asterisk indicate unspecific band. **e**, mRNA-1273 poly(A) lengths distribution for HEK293T cells with tetracycline-induced depletion of CNOT1 (CNOT1 KD) or controls (WT, cultured without tetracycline). Reads are divided into ones with (blue) and without (red) mΨCmΨAG sequence. Lower panel shows the relative abundance of each group. P.values calculated using Wilcoxon test, p. value adjustment with Benjamini-Hochberg method.

To verify the role of deadenylation in mRNA-1273 decay, we constructed a HEK293T line with tetracycline-inducible depletion of the scaffold subunit (CNOT1) of the primary deadenylase complex CCR4-NOT. CNOT1 depletion increased global poly(A) lengths of endogenous mRNAs (ED Fig. 2f) and also decreased poly(A) tail shortening in mRNA-1273, compared to the uninduced state (**Fig. 2e**). We only observed significant changes in poly(A) length in reads lacking the terminal mΨCmΨAG pentamer. The ratio of transcripts lacking the pentamer remained the same in both conditions, implying that CCR4-NOT is not involved in pentamer removal. These observations indicate that, as previously suggested based on the analysis of endogenous transcripts, deadenylation limits the lifetime of therapeutic mRNAs in model cell lines.

### Re-adenylation of mRNA-1273 *in vivo*

The exact fate of mRNA therapeutics depends on many factors, which are hard to recapitulate in model cell lines. The micro-environment at the site of RNA delivery and the gene expression patterns specific to cells taking up the mRNA may influence its stability and, thus, the therapeutic outcome. Hence, we intramuscularly delivered mRNA-1273 via an *in vivo* mouse model to study vaccine mRNA metabolism.

We isolated RNA from the tissues at the injection sites 2, 8, and 24 h post-immunization (**Fig. 3a**) and observed that vaccine mRNA quickly diminishes from the injection site and is barely detectable after 24 h (**Fig. 3b**). eDRS was conducted for samples where a sufficient amount of vaccine mRNA was detected (as revealed by qPCR). Of interest, the majority of mRNA-1273 poly(A) tails contained mΨCmΨAG 24 h after injection, though mild deadenylation of processed tails was noticeable at all analyzed time points (**Fig. 3c**). To our surprise, a substantial fraction of mRNA-1273 reads had increased poly(A) length, reaching up to 150-200As. All of these reads lacked the terminal pentamer, suggesting that re-adenylation of vaccine RNA occurs after its removal. At this juncture, most of the injected vaccine RNA would have been taken up by immune cells and transferred to draining lymph nodes^6^. Unfortunately, due to the overall low prevalence of mRNA-1273 in lymph nodes, compared to injection sites^6^, we could not analyze similar poly(A) tail extension dynamics there.

**Figure 3.**
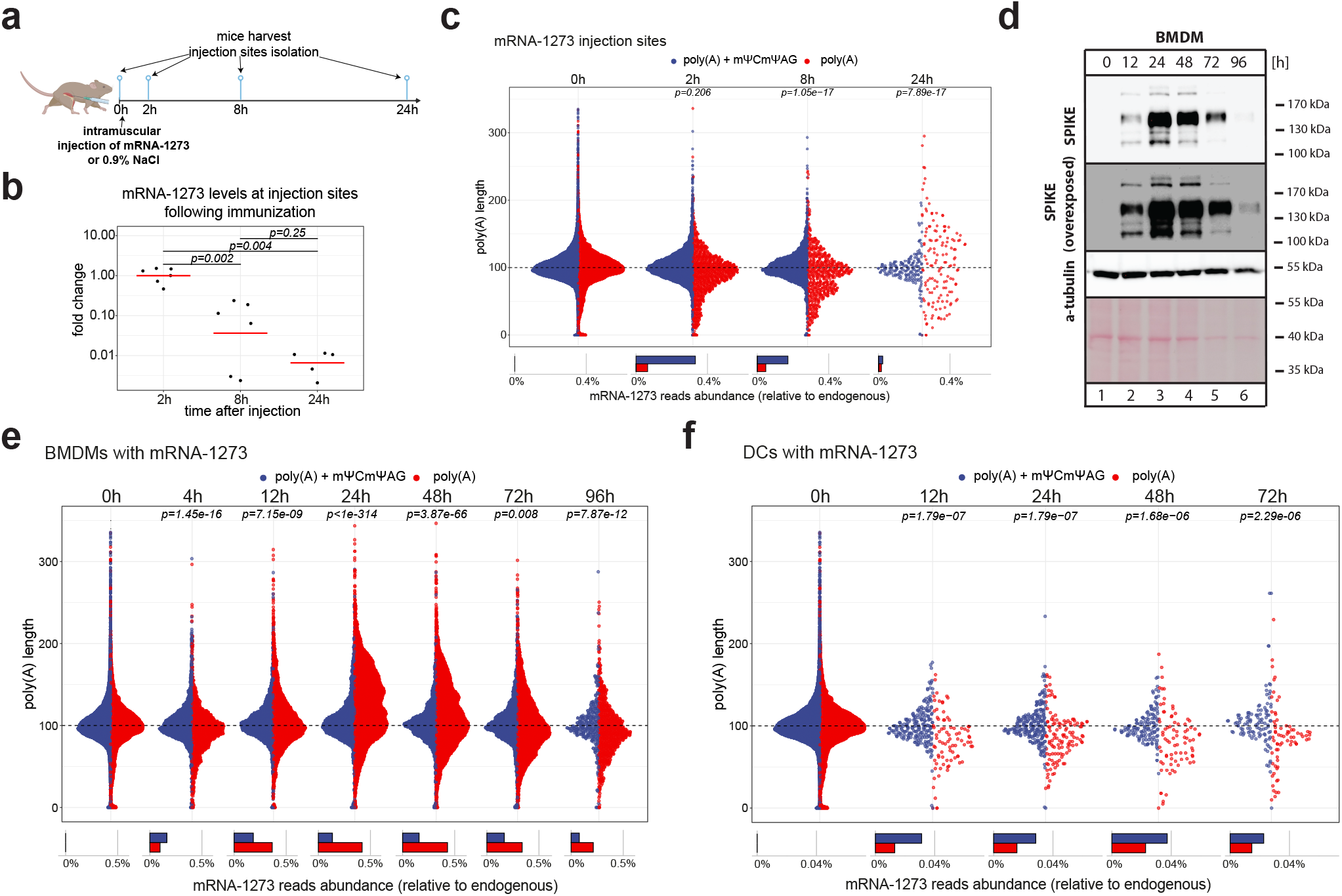
mRNA-1273 poly(A) tail extension after mΨCmΨAG removal *in vivo* and in macrophages. **a**, Schematic representation of the vaccination experiment **b**, Levels of mRNA-1273 at injection sites measured with qPCR 2, 8 and 24 h after injection of mRNA-1273. All values are shown as fold change relative to 2 h timepoint. Red lines represent mean values. **c**, mRNA-1273 poly(A) lengths distribution at mRNA-1273 injection sites up to 24 h post-injection. Average distributions for 2-3 replicates are shown. Reads are divided into ones with (blue) and without (red) mΨCmΨAG sequence. Lower panel shows the relative abundance of each group. P.values calculated using Wilcoxon test, p. value adjustment with Benjamini-Hochberg method. **d, E**xpression of mRNA-1273 in BMDMs up to 96 h after mRNA-1273 delivery. Western Blot on spike protein, α-tubulin as control. Asterisk indicate unspecific band. **e-f**, mRNA-1273 poly(A) lengths distribution in BMDMs (**e**) and DCs (**f**) treated with vaccine for up to 72 h (DCs) or 96 h (BMDMs). Average distributions for 2 (DCs) or 3 (BMDMs) replicates are shown. Reads are divided into ones with (blue) and without (red) mΨCmΨAG sequence. Lower panel shows the relative abundance of each group. P.values calculated using Wilcoxon test, p. value adjustment with Benjamini-Hochberg method.

To see whether mRNA-1273 re-adenylation occurs in monocyte/macrophages and DCs, which are reported to be the major cell type taking up BNT162b2^5^, we analyzed vaccine metabolism using eDRS in *in vitro* cultures of bone marrow-derived macrophages (BMDMs) and bone marrow-derived dendritic cells (BMDCs, further termed DCs). Both types of cells were efficiently transfected mRNA-1273 LNPs without visible toxic effects (ED Fig. 3a), allowing antigen translation for up to 72 h (**Fig. 3d**). Vaccine mRNA was stable and easily detectable for at least 72 h, with around 25% of molecules having an intact 3’end after 4 h (**Fig. 3ef**, ED Fig. 3bc). We observed that 24 h after transfection, the poly(A) tails of vaccine mRNAs were elongated by ~20As on average, reaching up to 200As in BMDMs (**Fig. 3e**, ED Fig. 3b). The mean poly(A) length returned to the initial ~100As 72 h after transfection. Notably, dynamic change in polyadenylation was observed only for tails lacking the 3’ terminal pentamer, and mRNA-1273 reads with an intact 3’ end had, on average, the same length of poly(A) at all analyzed time points. Re-adenylation was not observed in DCs (**Fig. 3f**, ED Fig. 3c).

In sum, our data show that mRNA-based vaccines can be stable and even undergo poly(A) tail elongation *in vivo*, a novel result that has not previously been reported or considered in the literature.

### Global transcriptomic responses upon mRNA-1273 administration and induction of TENT5 poly(A) polymerases

Gene expression analysis of RNA samples isolated from injection sites revealed complex transcriptomic responses following vaccination. Changes are already visible 8 h post-vaccination, with 1,129 genes downregulated and 2,502 genes upregulated at 24 h, clustered into 3 groups (**Fig. 4a**, Supplementary Table 1). Examination of upregulated genes (clusters 2 and 3) revealed significant enrichment in functions related to immune response and cell activation (**Fig. 4a**, ED Figure 4a, Supplementary Table 2). In addition, the expression of genes related to basic metabolic processes was decreased (**Fig. 4a**, cluster 1, ED Fig. 4a, Supplementary Table 1, Supplementary Table 2).

**Figure 4.**
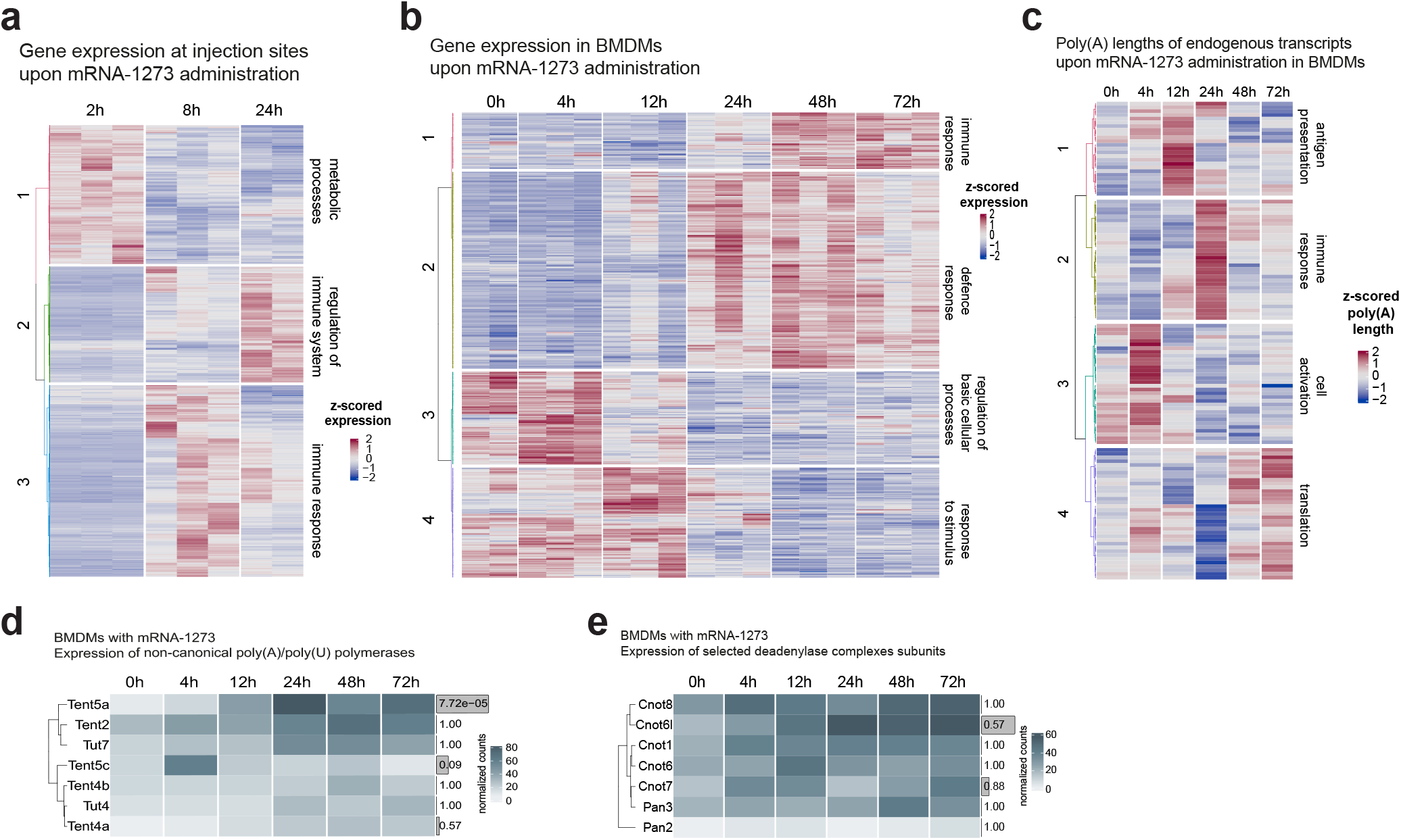
Innate immune response and induction of TENT5A poly(A) polymerase after vaccine administration. **a-b**, Genes with changed expression at injection sites (**a**) or in BMDMs (**b**) upon mRNA-1273 administration. Expression values were z-score normalized. Only genes with a significant change in expression (LRT test p<0.05) are shown. Clustering done using hclust, with ward.D method. On the right, descriptions on major functionalities enriched in given cluster are shown. N=3656 (a) or 584 (b). **c**, Transcripts with changed poly(A) lengths (revealed with Kruskal Wallis test) in the time course (up to 72 h) upon mRNA-1273 administration. Z-score normalized poly(A) lengths are represented in a colour-scale. Clustering done using hclust, with ward.D method. On the right, descriptions on major functionalities enriched in given cluster are shown. N=124. **d-e**, Expression of non-canonical poly(A)/poly(U) polymerases (**d**) or deadenylase complexes subunits (**e**) in BMDMs treated with mRNA-1273 for up to 72 h. Gene names are indicated on the left, on the right adjusted p.values from the DESeq2 LRT test are shown. Height of the bar indicates the expression change significance. Expression is shown as DESeq2-normalized counts.

Patterns of gene expression following mRNA-1273 administration in BMDM cultures were similar to those of transcriptomic responses at injection sites. 584 transcripts changed their expression, with the most pronounced transcriptome reconstruction observed 24 h after vaccine administration and persistent till the end of the experiment (72 h) (**Fig. 4b**, Supplementary Table 3). Those transcripts were clustered into 4 groups (**Fig. 4b**), two of which contained genes upregulated after 24 h and mainly associated with antigen presentation (MHC components in cluster 1), innate immunity (*ApoE, Lyz2, CtsH, CtsS*) and interferon induction (*Stat1, Mx1*, and multiple other Ifi transcripts) (**Fig 4b**, ED Fig. 4b, Supplementary Table 3, Supplementary Table 4). These changes resemble responses previously reported for the BioNTech mRNA anti-COVID19 vaccine (BNT162b2)^5^.

Intriguingly, in contrast to mRNA-1273 administration in BMDM cultures, almost no global changes in transcription were observed in DCs, where only certain innate immunity effectors (*Ctsb, Ctsd*) and serum complement element C1q (*C1qb*) were induced 72 h post administration (ED Fig. 4c, Supplementary Table 5).

As the vaccine mRNA is not expected to enter the nucleus, there should be a specific cytoplasmic mechanism causing the extension of vaccine poly(A) tails after delivery to macrophages. Whole-transcriptome poly(A) profiling revealed that the change in poly(A) length is not specific to mRNA-1273: the poly(A) tails of 124 endogenous transcripts were significantly changed after treatment with mRNA-1273 (Kruskall-Wallis test, p<0.001), with immune-response-associated transcripts exhibiting the most pronounced tail extension (average: 20As at 24 h, **Fig. 4c**; ED Fig. 4d, clusters 1 and 2; Supplementary Table 6, Supplementary Table 7). They included MHC components (*H2-K1, H2-D1, H2-T23, B2m*), serum complement components (*C1qb, C1qc*), lysosomal proteins (*Lamp1, Laptm5*), and innate immune response genes (*Ctsb, Ctsd, ApoE, Lyz2, Cst3, Ctss*). Remarkably, in our previous study on BMDMs activated with LPS^26^, a similar set of transcripts was revealed to be substrates of TENT5A/TENT5C non-canonical poly(A) polymerases. Therefore, we examined gene expression changes of all known non-canonical poly(A)/poly(U) polymerases and found that *Tent5a* is the only polymerase that is significantly induced at vaccine injection sites and after vaccine treatment in BMDMs (**Fig. 4d,** ED Fig. 4e). *Tent5c* is also expressed but at a much lower level. At the same time, no change in the expression of subunits of deadenylase complexes was observed (**Fig. 4e**). In contrast to BMDMs, both *Tent5a* and *Tent5c* were barely expressed in DCs. These experiments suggest a possible role for TENT5A and, to a lesser extent TENT5C, in the re-adenylation of mRNA-1273 that is observed in BMDMs, but not in DCs.

### mRNA-1273 polyadenylation by TENT5s is essential for the efficient production of spike protein and immune response

To evaluate the role of TENT5 proteins in vaccine mRNA re-adenylation, we studied mRNA-1273 metabolism in BMDMs devoid of TENT5A and TENT5C (double-knockout *Tent5a^Flox/Flox^/Tent5c*^-/-^)^26^. Double-knockout BMDM cells were treated with mRNA-1273 for up to 72 h, RNA was isolated and subjected to eDRS. In comparison to BMDM WT cells, we observed relatively fast mRNA-1273 decay and deadenylation in the absence of TENT5A and 5C (**Fig. 5a**, SD Fig. 5a), with mean poly(A) tails shortened to 80As after 24 h, despite the high ratio of RNAs lacking the 3’ pentamer. This indicates a critical role of TENT5A/C proteins in the regulation of mRNA-1273. Indeed, in *Tent5a^Flox/Flox^/Tent5c^-/-^* BMDMs stability of mRNA-1273 is drastically reduced (**Fig. 5c**).

**Figure 5.**
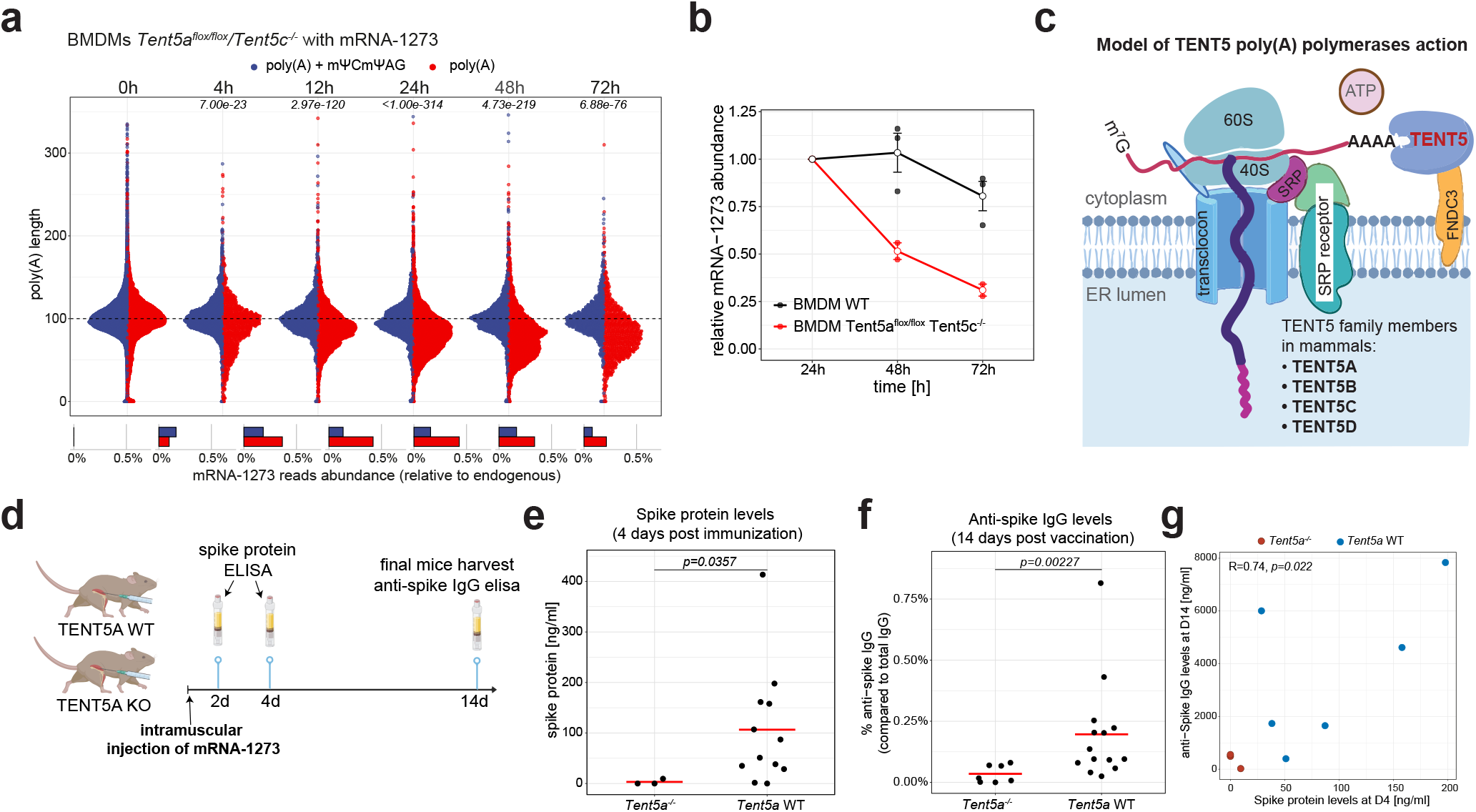
TENT5 poly(A) polymerases re-adenylate mRNA-1273, enhancing antigen production and immune response. **a**, mRNA-1273 poly(A) length distribution in BMDMs (*Tent5a^flox/flox^/Tent5c^-/-^*) treated with vaccine for up to 72 h. Average distributions for 2 replicates are shown. Reads are divided into ones with (blue) and without (red) mΨCmΨAG sequence. Lower panel shows the relative abundance of each group. P.values calculated using Wilcoxon test, p. value adjustment with Benjamini-Hochberg method. **b**, Stability of mRNA-1273 in WT (black) or *Tent5a^flox/flox^/Tent5c^-/-^* (red) BMDMs (based on eDRS, refer to Fig. 3e and Fig. 5a). Abundance of mRNA-1273 normalized to 24 h timepoint. Standard errors are shown as error bars. **c**, Model of action of TENT5 poly(A) polymerases, which target transcripts encoding proteins translated on the endoplasmic reticulum (ER). **d**, Overview of the immunization experiment. **e**, Spike protein levels in serum of immunized mice, WT and *Tent5a^-/-^*, measured 4 days after immunization. Red lines represent mean values. **f**, Anti-spike IgG levels in serum of immunized mice, WT and *Tent5a^-/-^*, measured 14 days after immunization. Values are shown in relation to overall IgG levels (refer to ED Fig. 5g). Red lines represent mean values. **g**, Correlation scatterplot showing relation between spike protein level at day 4 post-immunization and anti-spike IgG levels at day 14 post-immunization.

TENT5A/C also affected endogenous transcripts as no extension of poly(A) tails following vaccination was observed for mRNAs noticed in WT (ED Fig. 5b, Supplementary Table 6), including innate immunity effector transcripts (clusters 1,2), previously observed in LPS-activated macrophages^26^. Importantly, dysfunction of TENT5-mediated cytoplasmic polyadenylation did not affect the transcriptional response, which was similar for both WT and *Tent5a^Flox/Flox^*/*Tent5c* BMDMs (ED Fig. 5c, Supplementary Table 8). Most mRNAs with poly(A) tails elongated after vaccination in WT BMDMs, but not *Tent5a^Flox/Flox^*/*Tent5c^-/-^* BMDMs, (ED Fig. 5b) encode extracellular proteins translated at the endoplasmic reticulum (ER), in line with previous findings regarding TENT5A/C substrates^26,27^. Notably, mRNA-1273 also encodes an ER-targeted protein, which, as we show, hijacks TENT5s to extend its poly(A) tail. To confirm the specificity of TENT5A to mRNAs encoding proteins traversing the ER, we co-transfected WT BMDMs with purified mRNA-1273 RNA and firefly luciferase-encoding mRNA reporter (mΨ containing *in vitro* transcribed mRNA, with ~75As tail). After 24 h, we collected RNA for eDRS. The elongation of poly(A) tails was observed for mRNA-1273, but not for the luciferase reporter, which is not translated on the ER (ED Fig. 5de). Curiously, the poly(A) tail elongation of mRNA-1723 was even more pronounced than in previous experiments, possibly because of increased availability of mRNA to the polyadenylation machinery, as compared to LNP-based delivery.

DCs are major antigen-presenting cells and are expected to play a major role in immunity development after vaccination. However, DCs express low levels of TENT5, raising the question of whether TENT5-mediated stabilization of vaccine mRNA in macrophages affects immune response. TENT5C is expressed in B cells, and was shown to regulate the humoral response by polyadenylation of immunoglobulin transcripts^28^, whereas TENT5A was described only in the context of innate immunity^26^. Thus, we immunized WT and *Tent5a*^-/-^ mice with mRNA-1273, and measured antigen and antibody production (**Fig. 5d**). Significantly lower spike (S) protein concentrations were observed in *Tent5a^-/-^* mouse serum, both at days 2 and 4 post-immunization, compared to WT (**Fig. 5e**). Further, the S protein concentration was significantly decreased on day 4, compared to day 2 (ED Fig. 5f). Thus, mRNA-1273 stabilization through TENT5A-mediated re-adenylation has a direct effect on antigen production, both the amount and the longitude. However, some effects on macrophage secretory capability caused by Tent5a KO can also contribute to decreased levels of the antigen. Importantly, the level of serum anti-spike IgG 14 days post-immunization was significantly lower in *Tent5a^-/-^* and in 3 of 4 mice was barely detectable (**Fig. 5f**), despite the overall higher level of IgGs in the sera compared to WT animals (ED Fig. 5g). These data indicate that impaired immune response in *Tent5a^-/-^* animals is directly related to diminished antigen translation and subsequent presentation. Moreover, since a good correlation exists between the S protein concentration in serum and spike-specific IgG in immunized mice (**Fig. 5g**), we postulate that secretion of the antigen by macrophages after intramuscular administration of mRNA vaccines plays a significant role in building the immune response.

Overall, our results indicate that mRNA vaccine metabolism varies across cell types and that this has a significant influence on immunization efficacy.

## Discussion

The Covid-19 pandemic accelerated vaccine technology development, and the timely introduction of mRNA vaccines has played an essential role in slowing the spread of the disease-causing virus, SARS-CoV-2. This report provides the first evidence that therapeutic mRNAs are modified in cells, such that their poly(A) tails are extended by the cellular machinery. We also demonstrate that cellular mRNA processing varies between cell types and tissues. Here, we identify TENT5A cytoplasmic poly(A) polymerase as responsible for vaccine mRNA re-adenylation, and we show that re-adenylation enhances the stability of mRNA therapeutics in macrophages and affects the magnitude of the humoral immune response. Finally, we implement a highly efficient methodology based on direct RNA sequencing (DRS) for the analysis of mRNA therapeutics. Together, our results offer an explanation for the efficacy of existing mRNA vaccines and may profoundly influence the development of next-generation mRNA therapeutics.

After administration, mRNA vaccines build immune responses in a complex way. Before COVID-19, efforts related to mRNA vaccine development were primarily focused on developing anti-cancer agents^11,29^. Although mRNA design was usually optimized in cell lines, the aim was to target dendritic cells that are assumed to produce and present the antigen, to build both humoral and cellular adaptive immune responses. Here, for the first time, we show that macrophages play a critical role and are responsible for antigen production. Not only are monocytes/macrophages the major cell populations that take up vaccine mRNA^5,9^, but these cells also express TENT5A, which re-adenylates mRNA-1273, thereby increasing its stability and protein output. In *Tent5a*^-/-^ mutant mice, a significant drop in antigen production is observed, coinciding with the impaired development of an antigen-specific humoral response. Thus, it can be expected that spike (S) proteins produced by macrophages, either at the site of vaccine inoculation or in the draining lymph nodes (where LNPs are transported and captured by macrophages), are endocytosed by LN resident DCs, resulting in induction of Th1 and Tfh cells, and follicular DCs, resulting in induction of humoral immune response.

Although the data presented were obtained using mRNA-1273, similarity in the mode of action of BNT162b2 indicates a likely role for TENT5A for this vaccine as well. Moreover, the fact that TENT5 substrates must be terminated with poly(A) sheds new light on the different efficacies of two anti-COVID19 mRNA vaccine prototypes developed by CureVac (CV). The original prototype, which failed clinical trials due to low efficacy, had a heavily engineered 3’ end: its poly(A)_64_ was followed by poly(C)_30_ and a histone stem-loop^16,30^. In contrast, the second generation prototype, CV2nCoV, which was more potent in building immune response, terminates with a poly(A) tail^16^ and thus would be amenable for TENT5 mediated re-adenylation.

Development of therapeutic mRNAs goes beyond vaccines, and intensive efforts are underway to use mRNA to treat many mendelian diseases or induce the production of therapeutic proteins. In these cases, the main target is the liver, as the transfection of hepatocytes is very efficient after intravenous delivery of LNP-encapsulated mRNAs. There are also attempts to target other organs/cell types. Here, we show that various cell types metabolize mRNA vaccines differently. Only those expressing high levels of TENT5s would be expected to stabilize mRNA through re-adenylation. Although the mechanism of substrate recognition by TENT5 is largely unknown, ER targeting is a prerequisite feature (**Fig. 5c**). Further mechanistic research will be needed to decipher substrate recognition of TENT5 poly(A) polymerases and to explore their physiological roles. However, it is clear that target cells expressing high levels of TENT5s will be desirable for therapeutic mRNAs encoding secreted proteins. Moreover, better knowledge about substrate recognition will allow for the rational design of mRNAs much more efficiently polyadenylated by TENT5 and, therefore, more stable. Other tissue-specific mechanisms of mRNA stabilization may also be uncovered in the future. All these observations emphasize that the design of therapeutic mRNAs needs to be tailored to the target cells and the target site of the protein product.

## Methods

### Mice lines and immunization

All mice lines were generated by the CRISPR/Cas9-based method in the Genome Engineering Unit (https://crisprmice.eu/) as described in ref. ^24,25,30^. A double knockout *Tent5a^Flox/Flox^/Tent5c^-/-^* was described previously^26^. Mice were bred in the animal house of the Faculty of Biology, University of Warsaw, and maintained under conventional conditions^24,25,30^ in open polypropylene cages filled with wood chip bedding enriched with nest material and paper tubes. Mice were fed at libutum with a standard laboratory diet (Labofeed B, Morawski). Humidity in the rooms was kept at 55 ± 10%, the temperature at 22 °C ± 2 °C, at least 15 air changes per hour, and the light regime set at 12 h/12 h (lights on from 6:00 to 18:00). Health monitoring was performed regularly at the IDEXX laboratory.

Discarded remnant vaccination material (Moderna mRNA-1273, Spikevax) was used within manufacturer’s guidelines for stability. Because this was not available for purchase and as only remnant (otherwise to be discarded) material could be used at the time, we obtained approval from the Polish Ministry of Health (MMI.454.1.2021.TM).

Mice were immunized by intramuscular (*Vastus lateralis* region) injection either with 25μl of 0.9% NaCl (controls) or 25μl of Moderna’s mRNA-1273 at 40 ng/μl concentration. Tissues were collected after 2, 8 and 24 hours post-injection and snap-frozen in liquid nitrogen. Serum was collected at day 2, 4 and 14 after immunization. All animal experiments were approved by the II Local Ethical Committee in Warsaw affiliated with the University of Warsaw, Faculty of Biology (approval numbers: WAW2/71/2021, WAW2/129/2021, WAW2/95/2022)) and were performed according to Polish Law (Act number 653 266/15.01.2015) and in agreement with the corresponding European Union directive.

### ELISA

The concentrations of spike protein, anti-spike IgG and total mouse IgG in mice serum were measured with commercially available ELISA kits: Sinobiological (#KIT40591), Eagle Bioscience (#KBVH015-14) and (#OGG11-K01) respectively. All reactions were conducted according to the manufacturer’s recommendation.

### Cell cultures

A549 and HEK293 Flp-In T-REx cell lines and its derivate’s were cultured in Dulbecco’s modified Eagle’s medium (DMEM; Gibco) supplemented with 10% FBS (Gibco) and penicillin/streptomycin (Sigma-Aldrich) at 37°C in a 5% CO2 atmosphere until 80% confluency. To produce the HEK293 Flp-In T-REx cells (R78007, Thermo Fisher Scientific) cell line with conditional knock-down of CNOT1 we designed a tri-miRNA construct (listed below; stem-loop sequences ATGGAAGAGCTTGGATTTGAT; CTCCCTCAATTCGCCAACTTA; AGGACTTGAAGGCCTTGTCAA), which was cloned at the BspTI (AflII)+ NotI restriction sites in a pKK-RNAi vector (pKK-BI16 nucCherry EGFP-TEV^31^), which is a derivative of the pcDNA5/FRT/TO vector. 1 mln HEK293 Flp-In T-REx cells were grown on DMEM High glucose (Gibco) supplemented with 10% fetal bovine serum on 6-well plates. To transfect cells 300 ng of the miCNOT1 bearing construct was mixed with 1 μg of pOG44 plasmid, 250 μl of OptiMEM media and supplemented with 2 μl TransIT-2020 Transfection Reagent (Mirus, MIR5400). The transfection mix was incubated 20 min at room temperature and then added to the cell culture for 24 h. Then cells were transferred to a 60 mm plate and cultured in a DMEM high glucose media supplemented with 40μg/ml hygromycin and 8 μg/ml blasticidin for the first 6-7 days and then the media was replaced with media of increasing antibiotic concentration to 50 μg/ml hygromycin and 10 μg/ml blasticidin. Cells were grown until single colonies appeared 4 weeks later. A control of cells non-transfected with the pKK-BI16 plasmid was included to ensure the specific selection of the cell line.

Expression of exogenous miRNA genes was induced by the addition of doxycycline (Thermo Fisher Scientific) at a final concentration of 100 ng/ml. Cell enumeration was performed by crystal violet (Sigma, C3886) staining described by Xin Chen Lab (UCSF).

### Murine bone marrow-derived macrophages (BMDM) cell cultures

The primary BMDM cell cultures were established from the bone marrow monocytes isolated from *Tent5a^Flox/Flox^/Tent5c^-/-^* and wild-type mice. Young adult mice (12-25 weeks old) were sacrificed by cervical dislocation, then femurs and tibias were isolated and bone marrow was harvested by centrifugation-based protocol^32,33^. Bone marrow cells were plated in IMDM medium (Thermo Fisher Scientific; 21980065) supplemented with 10% FBS (Gibco), 100 U/ml penicillin/0.1 mg/ml streptomycin solution (Sigma-Aldrich), and 10 ng/ml macrophage colony-stimulating factor (M-CSF, Preprotech; 315-02) and cultured at 37 °C in 5% CO2. For conditional *Tent5a* gene targeting, BMDM cells were transduced on the 8th day after isolation with lentivirus carrying Cre recombinase (pCAG-Cre-IRES2-GFP). The lentivirus production, cell transduction and genotyping were performed as described previously^26,28^. Cells were used for experiments on the 14th day after isolation.

### Murine bone marrow-derived dendritic cells (BMDCs) isolation and culture

Bone marrow was flushed with cold PBS from the femurs of 6-week-old female C57BL/6 mice. RBCs were lysed using ACK (Ammonium-Chloride Potassium) Lysing Buffer (ThermoFisher Scientific) according to the manufacturer’s protocol. All the remaining cells were suspended at 1 x 10^6^/ml density in the culture medium [RPMI 1640 medium (Sigma-Aldrich) supplemented with heat-inactivated 10% (v/v) fetal bovine serum (HyClone), 2 mM l-glutamine (Sigma-Aldrich), 100 U/ml penicillin and 100 μg/ml streptomycin (both from Sigma-Aldrich)], plated at ø10 cm non tissue culture-treated Petri dishes (Sarstedt) and placed at 37 °C in the atmosphere of 5% CO_2_ in the air. Differentiation towards DCs was induced with 200 ng/ ml recombinant murine FLT3L (Peprotech). On day 7^th^ 600 pg/ml of recombinant murine GM-CSF (Peprotech; 315-03) was added for 4-day incubation. On day 11^th^ loosely adherent and floating cells were collected for subsequent experiments.

### mRNA-1273 administration to in vitro cultured cells

The 0.5-1M cells were seeded day before on 6-well plate in media as described above. 5 μl of mRNA-1273 (200 ng/μl) were diluted in 150 μl of Opti MEM media (Thermo Fisher Scientific), at room temperature and after 10 min. added to the cells in drop-wise manner and gently mixed. Cells were harvested at time points specified for the individual experiments.

The mRNA transfections, using purified mRNA-1273 and FLuc spike-in, were carried out with Lipofectamine MessengerMAX (Thermo Fisher Scientific), according to manufacturer’s instructions. Cells were harvested 24 h after transfection for subsequent analyses.

### General molecular biology techniques

#### Western blots

An equal number of cells were lysed in PBS supplemented with 0.1% NP40, protease inhibitors and viscolase (final concentration 0.1 U/ml; A&A Biotechnology, 1010-100) for 30 min at 37 °C with shaking 1200 rpm, then 3x SDS Sample buffer (187.5 mM Tris-HCl pH 6.8, 6% SDS, 150 mM DTT, 0.02% Bromophenol blue, 30% glycerol, 3% 2-Mercaptoethanol) was added and samples were boiled for 10 min. Samples were resolved on 12–15% SDS–PAGE gels and then proteins were wet transferred to Protran nitrocellulose membranes (GE Healthcare) at 400 mA at 4 °C for 1.5 h in 1x Transfer buffer (25 mM Tris base, 192 mM glycine, 20% methanol (v/v)). Next, the proteins were visualized by staining with 0.3% w/v Ponceau S in 3% v/v acetic acid and digitalized. Membranes were blocked by incubation in 5% milk in TBST buffer for 1 h followed by overnight incubation at 4 °C with specific primary antibodies (listed in Table 1) diluted 1:2500 (spike protein, firefly luciferase, and PDI) or 1:5000 (α-tubulin, β-actin, and GAPDH). Membranes were washed three times in TBST buffer, 10 min each, incubated with HRP-conjugated secondary antibodies: anti-mouse (Millipore, 401215) diluted 1:5000 and anti-rabbit (Millipore, 401393) diluted 1:5000, for 2 h at room temperature. Membranes were washed three times in TBST buffer and proteins were visualized using ChemiDoc System (BioRad) or X-ray films. Unprocessed scans of selected major blots and gels are shown in the Extended Data Fig. 6.

**Table 1.** Antibodies used in this study

| Antibodies | Manufacturer and catalog number |
| --- | --- |
| Anti-SARS-CoV-2 Spike Glycoprotein S1 antibody | Abcam, ab275759 |
| Anti-Firefly Luciferase | Abcam; ab21176 |
| Anti-GAPDH | Novus Biologicals, NB300-327 |
| Anti-Tubulin | mAb (DM1A), CP06 |
| Anti-Actin | Cell Signaling; 2146 |
| Anti-PDI (C81H6) | Cell Signaling; 3501 |
| Goat Anti-Mouse IgG, H&L Chain Specific Peroxidase Conjugate | Millipore Cat# 401215 |
| Goat Anti-Rabbit IgG, H&L Chain Antibody, Peroxidase Conjugated | Millipore Cat # 401393 |

Antibodies used in this study are listed in Table 1 below

#### RNA isolations

Total RNA was isolated from cells or vaccine samples with TRIzol reagent or TRIzol™LS Reagent (both from Thermo Fisher Scientific), respectively, according to the manufacturer’s instructions, dissolved in nuclease-free water and stored at −20 °C (short term) or −80 °C (long term). RNA from frozen muscles and lymph nodes were isolated by tissue homogenization in TRI-reagent (Sigma, T9424) pre-heated to 60 °C, using Omni Tissue Homogenizer (TH) equipped with 7 x 115 mm Saw Tooth (Fine) Generator Probe, homogenous mixtures were further processed according to the manufacturer’s instructions. For RT-qPCR, RNA-seq and DRS library preparation, the RNA was treated with TURBO DNase (Thermo Fisher Scientific; AM1907; or Invitrogen™, AM2238). To assess the integrity of the used material, each RNA sample after DNAse treatment was analyzed with Agilent 2200 TapeStation system, using Agilent High Senitivity RNA ScreenTape (Agilent, 5067-5579).

### mRNA-1273 quantification

cDNA was synthesized using 500 ng of DNAsed total RNA as template with SuperScript™ III Reverse Transcriptase (Invitrogen™, 18080093) according to manufacturer’s instructions. Final cDNA concentration was kept at 2,5 ng/ul (converted from total RNA).

For assessment of Moderna’s mRNA-1273 concentration in mouse tissues, custom made TaqMan probe was used together with TaqMan™ Gene Expression Master Mix (Applied Biosystems™, 4369016). Pre-designed gene expression assay for beta-Actin (Applied Biosystems™, Mm01205647_g1) was used for normalization. The reaction mix was contained in 10 μl total volume with 1x concentration of master mix and gene expression assay, 5 ng cDNA was used per reaction (converted from total RNA). Thermal cycling program for TaqMan™ Gene Expression Master Mix was used as instructed by the manufacturer.

For relative gene expression estimation of *Tent5a* and *Tent5c* at injection sites, and mRNA-1273 levels in HEK293T and A549 cells with varying amounts of vaccine, we used Platinum™ SYBR™ Green qPCR SuperMix-UDG (Invitrogen™, 11733046) following the general protocol for ABI instruments recommended by the manufacturer.

QuantStudio™ 5 Real-Time PCR System, 384-well (Applied Biosystems™, A28140) was used in case of all RT-qPCR analyses.

#### Custom TaqMan probe design

Primer blast algorithm was used to find a unique amplicon for mRNA-1273 within mouse transcriptome (Refseq mRNA). Within the amplified region, a 15-mer of optimal GC content (53,3%) was selected as target site for probe hybridization. We selected FAM dye and MGB-NFQ quencher as 5’ and 3’ modifications for the probe. The probe was synthesized by ThermoFisher’s Custom TaqMan^®^ Probes service. In order to test the specificity and sensitivity of designed assay we performed a serial dilution experiment in which we spiked 500 ng of DNAsed murine total RNA with 50 ng of DNAsed mRNA-1273 and performed cDNA synthesis. Then we performed 10x serial dilutions of said mix with unspiked murine cDNA of the same concentration. RT-qPCR analysis has shown that the designed assay is specific and can detect up to 10 ag of mRNA-1273 per ng of total RNA which is equivalent of roughly 25 molecules.

Primers and probe sequences as well as working concentrations were as listed in Table 2 below:

**Table 2.**
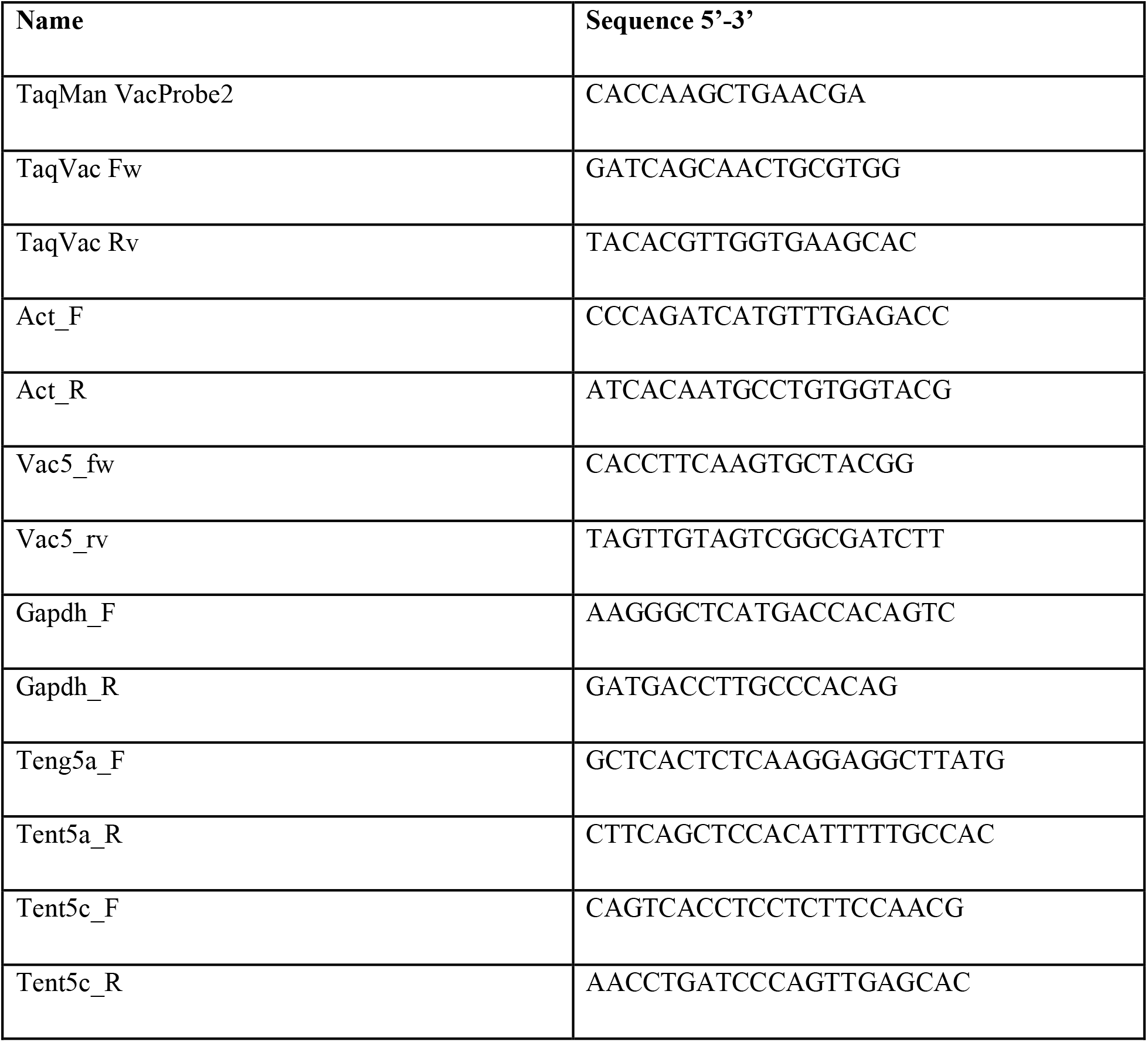
Primers and probe sequences

### 3’RACE-seq

To examine 3’UTR and terminal sequence of the mRNA-1273, RNA was freshly isolated from the vaccine sample vial. Then 1 μg of vaccine mRNA was ligated to 20 pmols RA3_7N adaptor: 5rApp/CTGACNNNNNNNTGGAATTCTCGGGTGCCAAGG/3ddC with 10U of T4 KQ227 RNA ligase 1 (#M0204S NEB) in the presence of 20U RNase OUT (#10777019, Thermo Fisher Scientific), 1x T4 RNA Ligase Reaction Buffer (#M0204S NEB), 1mM ATP, 20% PEG8000 in total 20 μl reaction volume at 25 °C for 4 h. Ligase was inactivated at 65 °C for 20 min. The ligation product was purified with 0.8x ratio KAPA Pure Beads and eluted with 15 μl RNase-free water, according to manufacturer protocol. The cleaned ligation product was subjected to reverse transcription with 40 pmol of Illumina index adapter: 5’CAAGCAGAAGACGGCATACGAGATATCAGTGTGACTGGAGTTCCTTGGCACCCGAGAATT CCA3’ with SuperScript III (#18080093, Thermo Fisher Scientific), 1x First Strand Buffer (#18080093, ThermoFisher), 0.25mM dNTP mix, 5mM DTT, 20U RNAse OUT (#10777019, Thermo Fisher Scientific). The reaction mix was incubated at 45 °C for 1 hour and 70 °C for 20 min in a thermocycler. Reverse transcription product was cleaned with 1x ratio KAPA Pure Beads and eluted with 19 μl RNase-free water. Prepared cDNA was diluted in 1:3 proportion. In the library amplification step, 0.5 μl of cDNA was mixed with 0.4 pmol of gene-specific starter 5’AATGATACGGCGACCACCGAGATCTACACGTTCAGAGTTCTACAGTCCGACGATCAGAAGGAGATCGATCGGCTG3’, 0.2 pmol of RP universal starter 5’CAAGCAGAAGACGGCATACGAGAT3’, 0.25mM dNTP, Phusion High-Fidelity DNA Polymerase (#F530S, Thermo Fisher Scientific), 1x Phusion HF Buffer (#F530S, ThermoFisher), 3% DMSO. For amplification, a standard Phusion program was used with 65 °C for annealing and 25 cycles. The 50 μl of PCR product was separated in 2.5% agarose gel. A band of 450bp was cut out from the gel and purified with Gel-OUT (#023-50, A&A Biotechnology) according to the kit protocol. The library was cleaned twice with 1.0x ratio of KAPA Pure Beads. TapeStation analysis of the sample was performed as quality control. The library was sequenced on Illumina NovaSeq 6000 sequencer.

### 3’-RACE-seq data analysis

RA37_N adapter sequence was trimmed from the R2 read (containing poly(A) tail) with cutadapt^34^ (options -g CCTTGGCACCCGAGAATTCCANNNNNNNGTCAG –discard-untrimmed). Then only reads containing poly(A) tail were identified with cutadapt (options -a TTTTTT --discard-untrimmed --fasta). Obtained sequences were reverse complemented using fastx_reverse_complement from the fastx toolkit (0.0.13) and loaded into R with BioStrings package. To get rid of unwanted trimming artifacts sequences with length between 0 and 10 were chosen, four A letters (representing poly(A) tail) were pasted before each sequence for visualization purposes, and sequence logo representing nucleotide composition of 3’ end of mRNA-1273 was produced with ggseqlogo package^34^.

### Preparation of standards with predefined poly(A) lengths

Spike-ins RNA were *in vitro* transcribed from a set of double-stranded DNA fragments. Templates for transcription were prepared in two consecutive PCR reactions. First, a desired fragment of Renilla luciferase from pRL5Box plasmid was amplified using RLuc_F1/R1 specific primers containing an overhang common for all primers utilized in the second round of PCR (Table 3). PCR products were verified through gel electrophoresis. Correct amplicons were used as templates in the second PCR reaction with RLuc_T7_F2 primer, hybridizing to the overhang sequence from RLuc_F1 and containing the T7 promoter, and backward primer RLuc_Ax_R2, hybridizing to the overhang sequence from RLuc_R1 oligo and introducing poly(A) tail of a defined length (from 10 to 120 As). Resulting PCR products were assessed and purified by gel electrophoresis. *In vitro* transcription reaction was performed at 37 °C for 1.5 h in a 50 μl reaction volume containing: 600 pmols T7 template, 10 μl of 5x transcription buffer (200mM Tris-HCl, 30mM MgCl_2_, 10mM spermidine, 50mM NaCl), 5 μl of rNTPs mix (20mM each), 5 μl of 100mM DTT, 0.5 μl of 1%Triton X-100, 80U Ribonuclease Inhibitor, 100U T7 RNA polymerase. Then, DNA template was removed with TURBO DNase (Ambion) for next 15min. Spike-ins RNA were phenol/chlorophorm extracted, precipitated, visually assessed by denaturing electrophoresis, purified on RNA purification beads and subjected as controls in DRS runs. In each DRS run, a mixture of spike-ins representing RNAs with poly(A) tail of defined length:A10/ A15/ A30/ A45/ A60/ A90/ A120 was included.

**Table 3.**
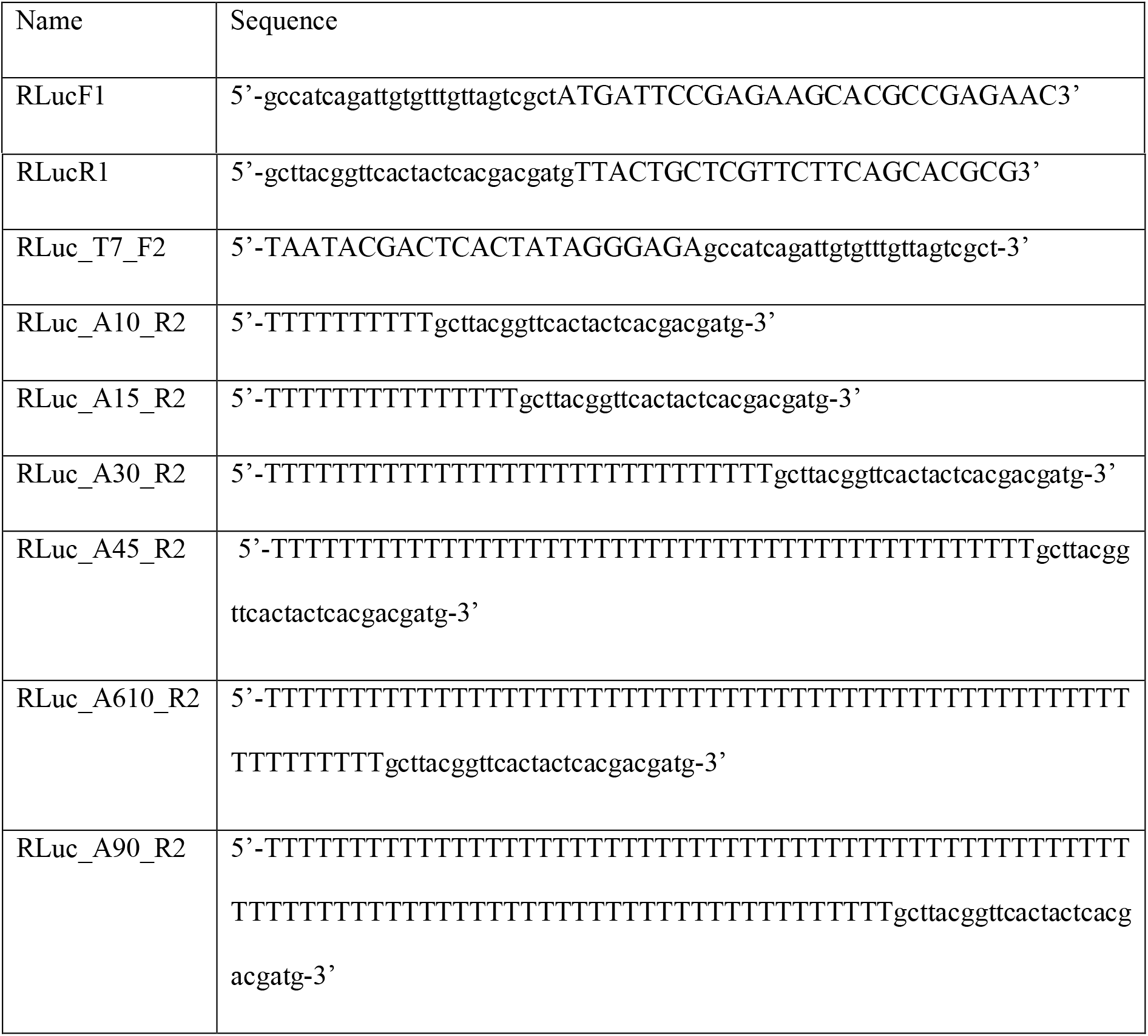
Primers used for spike-ins preparation

### FLuc mRNA synthesis

The Firefly Luciferase gene (FLuc) was purchased from Invitrogen (Thermo Fisher Scientific) with restriction sites for the AdeI and BamHI endonucleases, and cloned into the plasmid vector pJet (Thermo Fisher Scientific). The mRNA was synthesized by *in vitro* transcription using a DNA template with T7 promoter Φ6.5 (TAATACGACTCACTATAGGG). The standard transcription reaction (100 μl) was carried out for 60 min at 37 °C. The transcription mixture contained 5mM concentrations of m1Ψ/ATP/CTP (Thermo Fisher Scientific), 4mM GTP (Thermo Fisher Scientific), and 10mM trinucleotide cap1 analog-m^7^GpppA_m_pG^36^. The reaction mixture was complemented with RNA polymerase buffer (Thermo), 20mM MgCl_2_, 1U/μL RiboLock RNase Inhibitor (Thermo Fisher Scientific), 0.002 U/μl Inorganic Pyrophosphatase (Thermo Fisher Scientific), T7 RNA polymerase (20 U/μl, Thermo Fisher Scientific) and 40 ng/μl linearized plasmid pJet_FLucA90 as a DNA template. After 60 min incubation DNA template was digested with 7U of DNase I (Thermo Fisher Scientific) for 30 min at 37 °C. To stop the reaction, 8 μl of 500mM aqueous Na2EDTA solution was added. The crude RNA was purified using POROS™ Oligo (dT)25 Affinity Resin (Thermo Fisher Scientific) according to the manufacturer’s protocol. The mRNA was condensed to 100 μl volume using the Amicon^®^ Ultra-4 centrifugal filter (Milipore) and finally purified using HPLC and the RNASep™ Prep column (ADS Biotec), using the following conditions: eluent A −100mM TEAAc, eluent B - 200mM TEAAc/ACN; 10–35% of eluent B in 32 min, precipitated as sodium salt (3M NaOAc pH 5.2, isopropanol/ 80% EtOH) and dissolved in water. The concentration of mRNA was determined using absorbance measurements. To check the purity of the mRNA preparations, 100 ng of mRNA was applied to 1% agarose gel and electrophoresis (20 min, 140 V) was performed.

### Inosine tailing and Nanopore DRS

2 μg of RNA isolated from mRNA-1273 vaccine sample was denatured in the presence of 20U RNAse OUT (#10777019, ThermoFisher) at 65 °C for 3 min and immediately placed on ice. I-tailing reaction was donewith 0.5mM ITP (inosine triphosphate), 1x NEB 2.0 buffer (#B7002, NEB) and 2U of poly(U) polymerase (#M0337S, NEB) at 37 °C for 45 min and terminated by snap-freeze in liquid nitrogen. I-tailed RNA was cleaned twice on KAPA Pure Beads (#7983298001, Roche) in a 1x ratio and used for DRS library preparation. I-tailed samples require ligation of special adaptor RTA_C10, which contains 10 cytosines at the 3’end: 5’ GAGGCGAGCGGTCAATTTTCCTAAGAGCAAGAAGAAGCCCCCCCCCCCC 3’. To make the I-tailing specific RTA adaptor we followed ONT Direct RNA Sequencing - Sequence-Specific protocol (Oxford Nanopore Technologies). 0.5 μg of I-tailed mRNA-1273 was taken to the first ligation step of DRS library preparation. Library preparation was performed as given in ONT Direct Sequencing protocol.

### Nanopore sequencing

Direct RNA sequencing was performed as described by Bilska et al., 2020^28^. For raw vaccine isolate, 0.5 μg of RNA was used for the library preparation. For RNA isolates obtained from cell cultures or mouse tissues, 3.5-5 μg of total mRNA was mixed with 50-200 ng oligo-(dT)_25_-enriched mRNA from *Saccharomyces cerevisiae* yeast and standards with predefined poly(A) lengths and processed with a Direct RNA Sequencing Kit (catalog no. SQK-RNA002, Oxford Nanopore Technologies) according to the manufacturer’s instructions. Sequencing was performed using R9.4 flow cells on a MinION device (ONT). Raw data were basecalled using Guppy (ONT). Raw sequencing data (fast5 files) were deposited at the European Nucleotide Archive (ENA, project PRJEB53190).

### Identification of mRNA-1273-originating reads using subsequence Dynamic Time Warping

For the comparisons of raw signals from nanopore sequencing, the DTAIDistance library^35^ (version 2.3.5) was used. Reference signal was obtained from one of the reads from the DRS run of crude mRNA-1273 material (read id: df408ab3-7418-4ee4-9a67-92743257b20a), which was giving good coverage of 3’end of mRNA-1273 reference. The 5000 data points, covering part of poly(A) tail (1000 data points) and 3’ end of transcript (4000 data points) were selected for further processing (ED Fig. 1b). Such fragment of raw current readout was smoothed using Savitzky-Golay filter (savgol_filter function from Scipy Python library), with window length set to 51 and polynomial order set to 3, and normalized using zscore function from stats python package. Raw sequencing data were read from fast5 files using ONT Fast5 API (version 4.0.0), first 20000 data points (which usually covers the 3’ end of sequenced RNA) were selected, then smoothed and normalized in the same way as reference signal. Then, the subsequence_alignment function from DTAIdistance library was used to find a region in raw data with the best match to the reference signal. As the output for each sequencing read the location of match and distance score, calculated using distance_fast function from DTAIDisatance library, was reported. The python script for all described operations is available at https://github.com/LRB-IIMCB/DTW_mRNA-1273

### Poly(A) lengths determination and statistical analysis

Basecalled nanopore reads were mapped to respective transcriptome reference using Minimap2 2.17 with options -k 14 -ax map-ont –secondary=no and processed with samtools 1.9 to filter out supplementary alignments and reads mapping to reverse strand (samtools view -b -F 2320). The poly(A) tail lengths for each read were estimated using the Nanopolish 0.13.2 polya function^25^.

For the analysis of mRNA-1273-originating reads the nanopolish polya algorithm was modified to (1) include unmapped reads, what allowed analysis of poly(A) lengths for sDTW-identified reads (2) detect mΨCmΨAG at the 3’ end of poly(A) tail and report its presence in the output.

For the detection of mΨCmΨAG the original segmentation algorithm was modified to include additional segment between adaptor and poly(A), with mixed Gaussian distribution emissions, which were manually estimated using MLE. As the result 4 additional columns are added to the nanopolish-polya output, containing localization of mΨCmΨAG in the raw signal, its length, summed poly(A) + mΨCmΨAG length, and information if mΨCmΨAG was detected. Detection of mΨCmΨAG is run with the nanopolish polya-moderna function. The source code of modified nanopolish is available at https://github.com/LRB-IIMCB/nanopolish_Moderna.

P values for each transcript were estimated using the Kruskal-Wallis test and adjusted for multiple comparisons using the Benjamini–Hochberg method. Transcripts were considered as having a significant change in poly(A) tail length, if the adjusted P value was < 0.05, and there were at least 20 supporting reads for each condition.

### Differential Expression analyses

Illumina RNA-seq reads were mapped to the mouse reference genome (GRCm38, ENSEMBL, release 94) using the STAR aligner (v 2.7.6a)^36^. Read counts were assigned to genes using featureCounts from Subread package (v. 2.0.1) with options -Q 10 -p -B -C -s 2 -g gene_id -t exon and respective annotation file (Gencode vM25). Multimappers and reads overlapping multiple features were not counted. Nanopore reads counts were derived from the nanopolish-polya output files, used for the poly(A) tail lengths analyses.

Differential expression analysis was performed with DESeq2 (v. 1.22) Bioconductor package^37^, using LikeLihood Ratio test for time-course experiments data. Genes with similar expression patterns were clustered with hierarchical clustering using ward.D method and Z-scored expression values for each gene. Heatmaps were drawn with the ComplexHeatmap package^38^. Gene Ontology enrichment analysis was done with g:Profiler^39^.

### Statistics and reproducibility

No statistical method was used to predetermine sample size. Statistical analysis was conducted on data from two or more biologically independent experimental replicates. Statistical analysis of quantitative data was performed using R environment unless otherwise stated. The statistical tests used in each instance are mentioned in the figure legends. All data were checked for normality using the Shapiro–Wilk test. Data are presented as scatter dot plots with mean values indicated and standard errors shown as the error bars, as indicated in the figure legends, and individual data points are shown. Most of the experiments were repeated at least twice, leading to comparable results with exception of mRNA-1273 treatment of A549 cells and viability assays, which were repeated once. Samples with clear technical failures during tissue harvesting, cells isolation, processing, or data collection were excluded from analyses.

## Data availability

Nanopore direct RNA sequences are deposited at the European Nucleotide Archive; accession number PRJEB53190. mRNA expression data are deposited in GEO. Raw data underlying figures are provided in ED Fig. 6 and Supplementary Datasets or are also available from the corresponding authors upon reasonable request.

## Code availability

Dynamic Time Warping script is available at https://github.com/LRB-IIMCB/DTW_mRNA-1273. Nanopolish-polya for identification of mΨCmΨAG is available at https://github.com/LRB-IIMCB/nanopolish_mRNA-1273.

## Acknowledgments

We thank Dziembowski lab members and IIMCB core facilities for help in selected experiments, Dr Małgorzata Piotrowska from the Medical Center of the Medical University of Warsaw for providing remnant vaccines and members of the Laboratory of RNA Biology, IIMCB, for help with vaccination experiments. *Tent5a-/-* and *Tent5a^Flox/Flox^/Tent5c^-/-^* mice were generated by the Genome Engineering Unit, IIMCB. Illumina sequencing was carried out by Genomics Core Facility, Centre of New Technologies, University of Warsaw (RRID:SCR_022718).

This work was supported by: the Virtual Research Institute, Polish Science Fund (to A.D, S.M, D.N., J.G., J.K. and J.J.); Foundation for Polish Science, financed by the European Union via the European Regional Development Fund (agreement no. TEAM/2016-1/3 to A/D) and National Science Center (SONATA BIS 10 2020/38/E/NZ2/00372 to S.M, OPUS 17 2019/33/B/NZ2/01773 to A.D., SONATA 16 2020/39/D/NZ2/0217 to A.T, and PRELUDIUM 19 2020/37/N/NZ2/02893 to A.B.). This research was supported by funding from the European Union Horizon 2020 research and innovation programme, under grant agreement no 810425.

## Author Information

These authors contributed equally: Olga Gewartowska, Michal Mazur, Wiktoria Orzel

## Contributions

A.D., S.M. conceived and designed the project. P.K. developed bioinformatic tools and analyzed all sequencing data. S.M. established the BMDM cultures, performed all cell line experiments, and isolated RNA for DRS. O.G. performed mice immunizations, serum collection, and ELISA measurements. W.O. carried out RNA isolation, DRS library preparation and sequencing, and 3’-RACE-seq experiment. M.M. did RNA isolation from injection sites and qPCRs. A.T. established the HEK293T CNOT1 KD cell line. K.M-K. prepared poly(A) length standards and participated in I-tailing experiment (together with W.O.). T.S. prepared the fluc spike-in. P. T., D.N, and J.G. provided the BMDC cultures. B.T. took part in immunization experiments. A.B. established the *Tent5a^Flox/Flox^/Tent5c^-/^* BMDMs cultures. S.J. did qPCRs on varying amounts of mRNA-1273 in HEK293T and A549 cells. J.K., J.G, K.M.K and S.M. provided feedback on the manuscript. A.D. and S.M supervised the work. P.K. and A.D wrote the manuscript, with contributions from other authors. All authors read and approved the manuscript.

## REFERENCES

1. Corbett, K. S. et al. SARS-CoV-2 mRNA Vaccine Design Enabled by Prototype Pathogen Preparedness. Nature 586, 567–571 (2020).

2. Sahin, U. et al. BNT162b2 vaccine induces neutralizing antibodies and poly-specific T cells in humans. Nature 595, 572–577 (2021).

3. Sahin, U., Karikó, K. & Türeci, Ö. mRNA-based therapeutics — developing a new class of drugs. Nat Rev Drug Discov 13, 759–780 (2014).

4. Szabó, G. T., Mahiny, A. J. & Vlatkovic, I. COVID-19 mRNA vaccines: Platforms and current developments. Molecular Therapy 30, 1850–1868 (2022).

5. Li, C. et al. Mechanisms of innate and adaptive immunity to the Pfizer-BioNTech BNT162b2 vaccine. Nat Immunol 1–13 (2022) doi:10.1038/s41590-022-01163-9.

6. Lindsay, K. E. et al. Visualization of early events in mRNA vaccine delivery in non-human primates via PET–CT and near-infrared imaging. Nat Biomed Eng 3, 371–380 (2019).

7. Ols, S. et al. Route of Vaccine Administration Alters Antigen Trafficking but Not Innate or Adaptive Immunity. Cell Reports 30, 3964–3971.e7 (2020).

8. Verbeke, R., Hogan, M. J., Loré, K. & Pardi, N. Innate immune mechanisms of mRNA vaccines. Immunity 55, 1993–2005 (2022).

9. Liang, F. et al. Efficient Targeting and Activation of Antigen-Presenting Cells In Vivo after Modified mRNA Vaccine Administration in Rhesus Macaques. Molecular Therapy 25, 2635–2647 (2017).

10. Diken, M. et al. Selective uptake of naked vaccine RNA by dendritic cells is driven by macropinocytosis and abrogated upon DC maturation. Gene Ther 18, 702–708 (2011).

11. Kranz, L. M. et al. Systemic RNA delivery to dendritic cells exploits antiviral defence for cancer immunotherapy. Nature 534, 396–401 (2016).

12. Karikó, K. et al. Incorporation of Pseudouridine Into mRNA Yields Superior Nonimmunogenic Vector With Increased Translational Capacity and Biological Stability. Molecular Therapy 16, 1833–1840 (2008).

13. Svitkin, Y. V. et al. N1-methyl-pseudouridine in mRNA enhances translation through eIF2α-dependent and independent mechanisms by increasing ribosome density. Nucleic Acids Research 45, 6023–6036 (2017).

14. Anderson, B. R. et al. Incorporation of pseudouridine into mRNA enhances translation by diminishing PKR activation. Nucleic Acids Research 38, 5884–5892 (2010).

15. Andries, O. et al. N1-methylpseudouridine-incorporated mRNA outperforms pseudouridine-incorporated mRNA by providing enhanced protein expression and reduced immunogenicity in mammalian cell lines and mice. Journal of Controlled Release 217, 337–344 (2015).

16. Gebre, M. S. et al. Optimization of non-coding regions for a non-modified mRNA COVID-19 vaccine. Nature 601, 410–414 (2022).

17. Lutz, J. et al. Unmodified mRNA in LNPs constitutes a competitive technology for prophylactic vaccines. npj Vaccines 2, 1–9 (2017).

18. Vogel, A. B. et al. BNT162b vaccines protect rhesus macaques from SARS-CoV-2. Nature 592, 283–289 (2021).

19. Corbett, K. S. et al. mRNA-1273 protects against SARS-CoV-2 beta infection in nonhuman primates. Nat Immunol 22, 1306–1315 (2021).

20. Xia, X. Detailed Dissection and Critical Evaluation of the Pfizer/BioNTech and Moderna mRNA Vaccines. Vaccines 9, (2021).

21. Gagne, M. et al. mRNA-1273 or mRNA-Omicron boost in vaccinated macaques elicits similar B cell expansion, neutralizing responses, and protection from Omicron. Cell 185, 1556–1571.e18 (2022).

22. Garalde, D. R. et al. Highly parallel direct RNA sequencing on an array of nanopores. Nature Methods 15, 201–206 (2018).

23. Fleming, A. M. & Burrows, C. J. Nanopore sequencing for N1-methylpseudouridine in RNA reveals sequence-dependent discrimination of the modified nucleotide triphosphate during transcription. 2022.06.03.494690 Preprint at https://doi.org/10.1101/2022.06.03.494690 (2022).

24. Kovaka, S., Fan, Y., Ni, B., Timp, W. & Schatz, M. C. Targeted nanopore sequencing by real-time mapping of raw electrical signal with UNCALLED. Nat Biotechnol 39, 431–441 (2021).

25. Workman, R. E. et al. Nanopore native RNA sequencing of a human poly(A) transcriptome. Nat Methods 1–9 (2019) doi:10.1038/s41592-019-0617-2.

26. Liudkovska, V. et al. TENT5 cytoplasmic noncanonical poly(A) polymerases regulate the innate immune response in animals. Science Advances 8, eadd9468 (2022).

27. Gewartowska, O. et al. Cytoplasmic polyadenylation by TENT5A is required for proper bone formation. Cell Reports 35, 109015 (2021).

28. Bilska, A. et al. Immunoglobulin expression and the humoral immune response is regulated by the non-canonical poly(A) polymerase TENT5C. Nature Communications 11, 2032 (2020).

29. Beck, J. D. et al. mRNA therapeutics in cancer immunotherapy. Molecular Cancer 20, 69 (2021).

30. Rauch, S. et al. mRNA-based SARS-CoV-2 vaccine candidate CVnCoV induces high levels of virus-neutralising antibodies and mediates protection in rodents. npj Vaccines 6, 1–9 (2021).

31. Mroczek, S. et al. The non-canonical poly(A) polymerase FAM46C acts as an onco-suppressor in multiple myeloma. Nature Communications 8, 619 (2017).

32. Amend, S. R., Valkenburg, K. C. & Pienta, K. J. Murine Hind Limb Long Bone Dissection and Bone Marrow Isolation. JoVE (Journal of Visualized Experiments) e53936 (2016) doi:10.3791/53936.

33. Bertović, I., Bura, A. & Jurak Begonja, A. Developmental differences of in vitro cultured murine bone marrow-and fetal liver-derived megakaryocytes. Platelets 33, 887–899 (2022).

34. Martin, M. Cutadapt removes adapter sequences from high-throughput sequencing reads. EMBnet.journal 17, 10–12 (2011).

35. Wagih, O. ggseqlogo: a versatile R package for drawing sequence logos. Bioinformatics 33, 3645–3647 (2017).

36. Sikorski, P. J. et al. The identity and methylation status of the first transcribed nucleotide in eukaryotic mRNA 5’ cap modulates protein expression in living cells. Nucleic Acids Research 48, 1607–1626 (2020).

37. Meert, W., Hendrickx, K., van Craenendonck, T. & Robberechts, P. DTAIDistance (Version v2.3.5). (2022) doi:10.5281/zenodo.5901139.

38. Dobin, A. et al. STAR: ultrafast universal RNA-seq aligner. Bioinformatics 29, 15–21 (2013).

39. Love, M. I., Huber, W. & Anders, S. Moderated estimation of fold change and dispersion for RNA-seq data with DESeq2. Genome Biology 15, 550 (2014).

40. Gu, Z., Eils, R. & Schlesner, M. Complex heatmaps reveal patterns and correlations in multidimensional genomic data. Bioinformatics 32, 2847–2849 (2016).

41. Raudvere, U. et al. g:Profiler: a web server for functional enrichment analysis and conversions of gene lists (2019 update). Nucleic Acids Research 47, W191–W198 (2019).

