## Extended Data Figures for "SARS-CoV-2 mRNA vaccine is re-adenylated *in vivo*, enhancing antigen production and immune response"

### Extended Data Figure 1

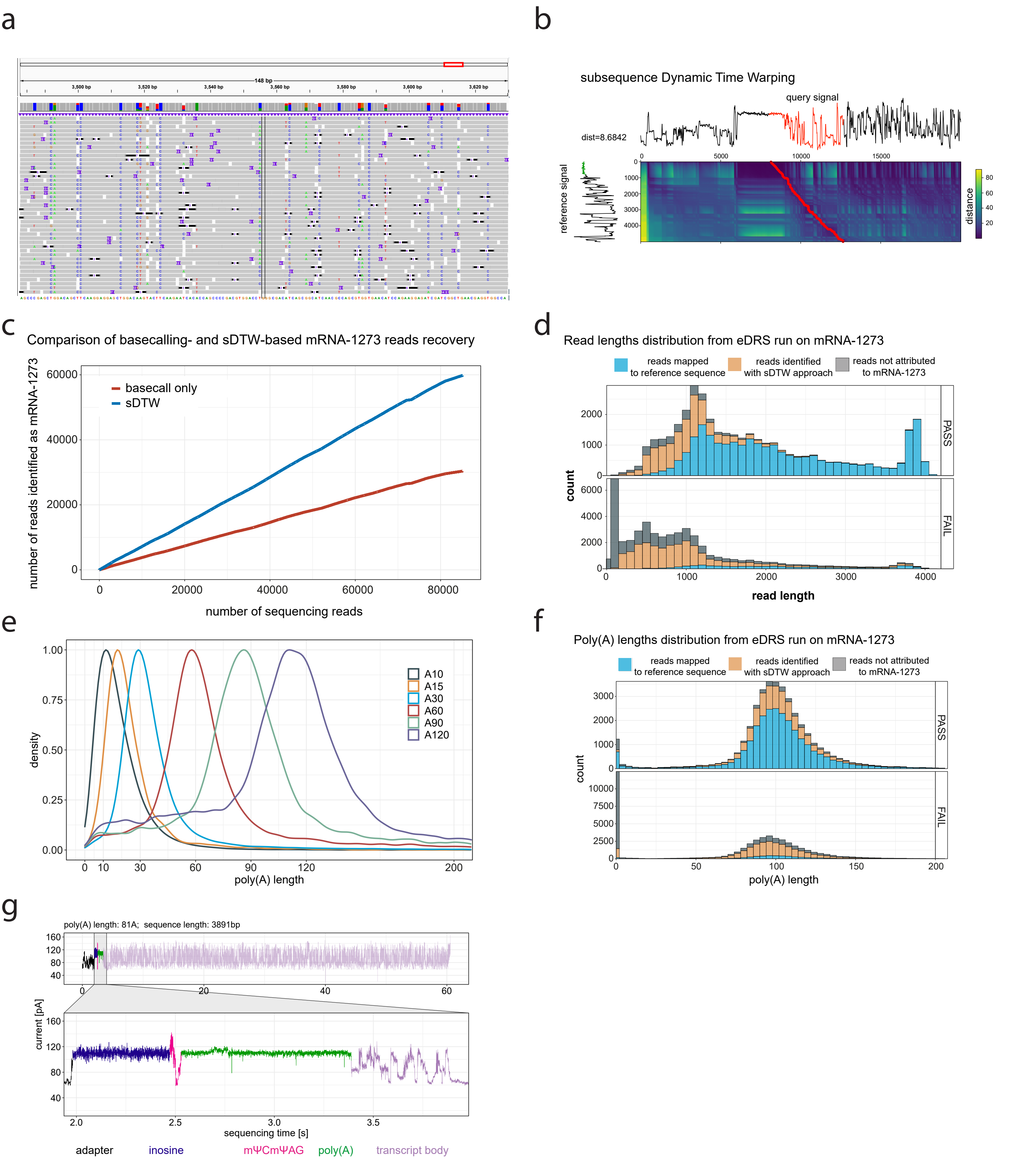

**Extended Data Figure 1. Enhanced Direct RNA Sequencing of mRNA-1273 (related to Figure 1).**

**a,** IGV snapshot of Guppy basecalled mRNA-1273 mapped to reference. Mismatches are visible in positions with mΨ incorporation.

**b,** Subsequence Dynamic Time Warping Approach (sDTW) approach. Reference signal (on the left, with the part of poly(A) tail marked in green) is compared with the query signal (on top), and a subsequence match location is found (marked in red). Heatmap shows the similarities between different parts of reference and query signals, used for drawing the warping path (in red).

**c,** Comparison of mRNA-1273 recovery using standard basecalling and DTW approach.

**d,** Distribution of read lengths from mRNA-1273 eDRS sequencing, for reads passing quality filter (mean qscore >=7, bottom panel) or failing (mean qscore <7, upper panel). In blue– reads basecalled using Guppy and aligned to the reference sequence. In yellow– reads identified as mRNA-1273 using DTW approach. In grey– other reads. Bin width = 100, stacked histogram.

**e**, Poly(A) lengths distribution of RNA spike-ins with predefined poly(A) lengths.

**f,** Distribution of poly(A) lengths for mRNA-1273 eDRS reads, for reads passing quality filter (mean qscore >=7, bottom panel) or failing (mean qscore <7, upper panel). In blue– reads basecalled using Guppy and aligned to reference sequence. In yellow– reads identified as mRNA-1273 using DTW approach. In grey– other reads. Bin width = 4, stacked histogram.

**g,** Representative raw eDRS signal from I-tailed mRNA-1273. Signal from poly(A) is zoomed in the bottom panel. In violet – transcript body, in green– poly(A) tail, in black – adaptor, in pink- mΨCmΨAG, in blue – poly(I) tail.

### Extended Data Figure 2

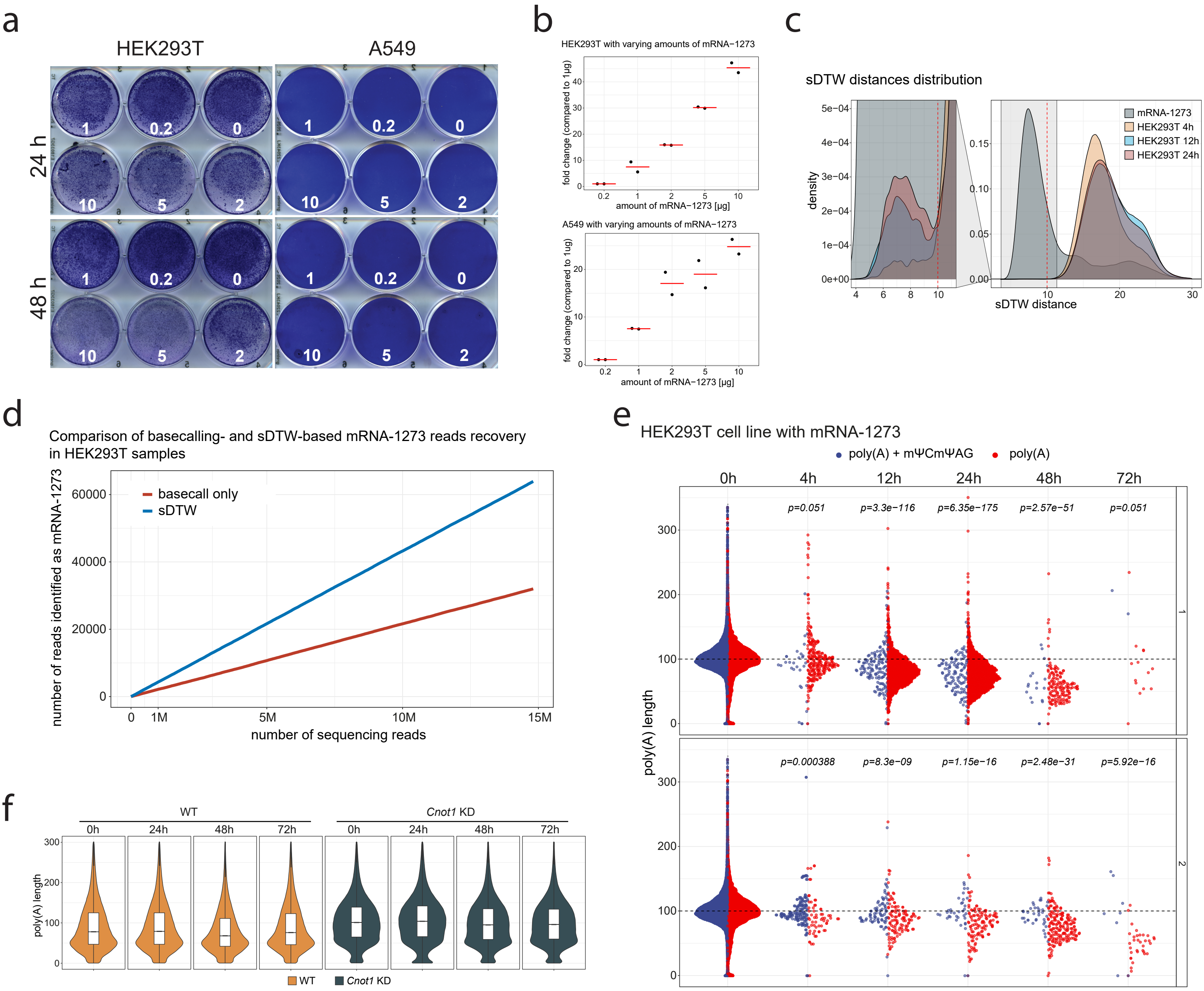

**Extended Data Figure 2. Stability of mRNA-1273 in model cell lines is determined by the deadenylation rate (related to Figure 2).**

**a-b,** Viability (**a**) and amount of vaccine RNA (**b**) in HEK293T or A549 cells incubated with 0-10 µg of mRNA-1273 per well of 6-well plate for 24 or 48 h assessed with crystal violet staining and qPCR, respectively. qPCR was performed on RNA isolated after 24 h of incubation (N=2).

**c**, Distribution of sDTW distances (compared to reference signal) of crude mRNA-1273 sample, and HEK293T samples incubated with mRNA-1273 (replicate 1, for clarity only 4, 12, and 24 h timepoints are shown). Distance threshold (<=10) for identification of reads as originating from mRNA-1273 indicated as red line.

**d,** Comparison of mRNA-1273 reads recovery from sequencing data of HEK293T + mRNA-1273 cells, using standard basecalling and sDTW approach.

**e,** mRNA-1273 poly(A) lengths distribution in HEK293T cells incubated with vaccine for up to 72 h, individual replicates shown. Reads are divided into ones with (blue) and without (red) mΨCmΨAG sequence. P.values calculated using Wilcoxon test, p. value adjustment with Benjamini-Hochberg method.

**f,** Distribution of poly(A) lengths of endogenous transcripts in HEK293T cells with, or without induction of CNOT1 depletion in right and left panels, respectively.

### Extended Data Figure 3

**a**

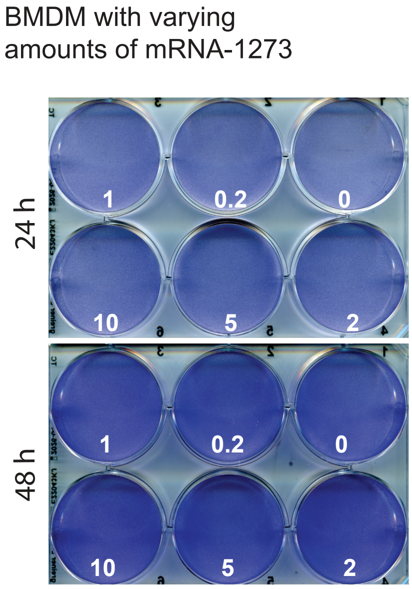


**b**

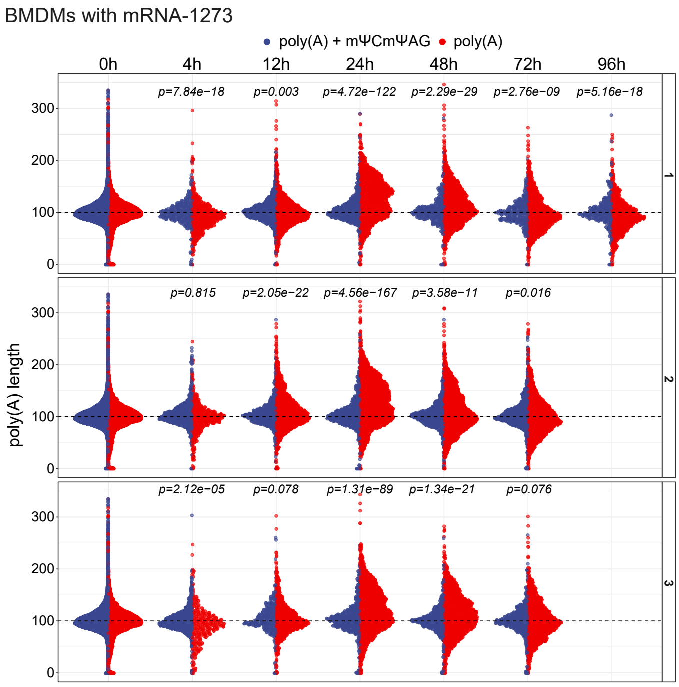


**c**

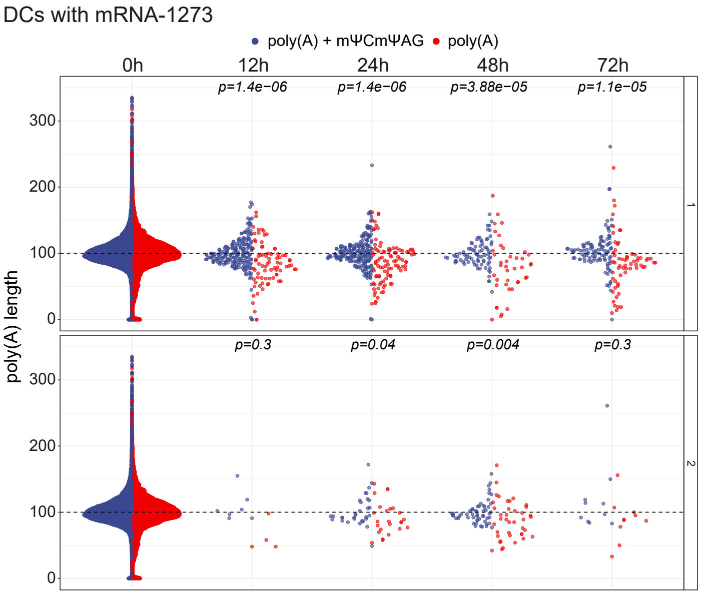

**Extended Data Figure 3. mRNA-1273 poly(A) tail extension after mΨCmΨAG removal *in vivo* and in macrophages (related to Figure 3).**

**a,** Viability of BMDMs incubated with 0-10 μg of mRNA-1273 per well of 6-well plate for 24 or 48 h, assessed with crystal violet staining.

**b-c,** mRNA-1273 poly(A) lengths distribution in BMDMs (**b**) and DCs (**c**) incubated with mRNA-1273 vaccine for up to 72h (DCs) or 96 h (BMDMs), Individual replicates are shown. Reads are divided into ones with (blue) and without (red) mΨCmΨAG sequence. P.values calculated using Wilcoxon test, p. value adjustment with Benjamini-Hochberg method.

### Extended Data Figure 4

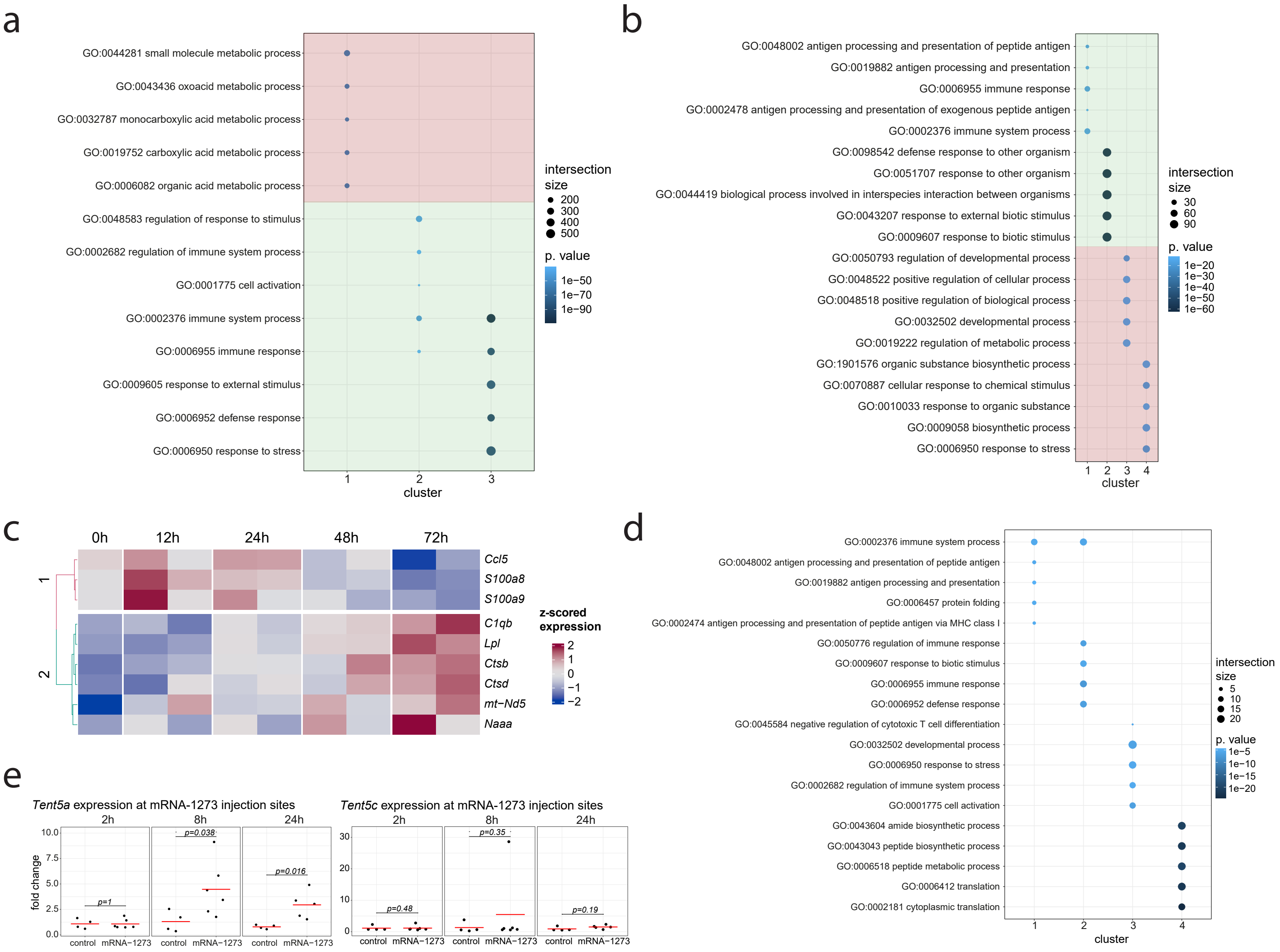

**Extended Data Figure 4. Innate immune response and induction of TENT5A poly(A) polymerase after vaccine administration (related to Figure 4).**

**a-b**, GO terms (category: Biological Process) enriched for transcripts in each of expression cluster identified for injection sites (**a**) or BMDMs (**b**) after mRNA-1273 administration. Respective clusters are visualized on Fig 4a and Fig 4b. GO terms were identified with G:profiler2.

**c,** Genes with changed expression in DCs upon mRNA-1273 administration. Expression values were z-score normalized. Only genes with significant change in expression (LRT test p.value<0.05) are shown. Clustering done using hclust, with ward.D method.

**d,** GO terms (category: Biological Process) enriched for transcripts in each of poly(A) length cluster identified for BMDMs after mRNA-1273 administration. Respective clusters are visualized on Fig 4c. GO terms were identified with G:profiler2.

**e,** Expression of *Tent5a* (left panel) or *Tent5c* (right panel) at mRNA-1273 injection sites measured with qPCR. P. values calculated with Wilcoxon test.

### Extended Data Figure 5

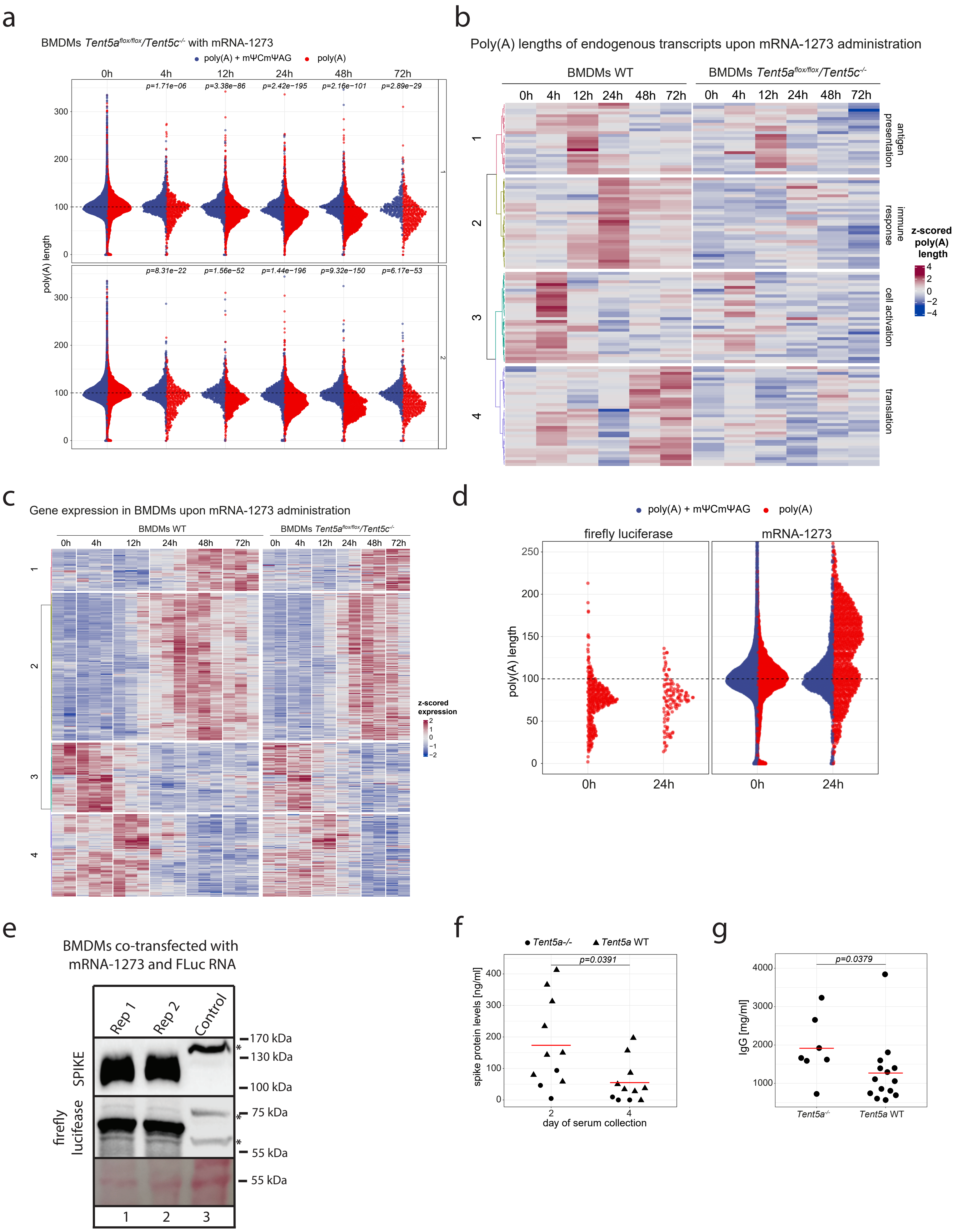

**Extended Data Figure 5. TENT5 poly(A) polymerases re-adenylate mRNA-1273, enhancing antigen production and immune response (related to Figure 5).**

**a,** mRNA-1273 poly(A) lengths distribution in BMDMs (*Tent5a*$^{flox/flox}$/*Tent5c*$^{-/-}$) treated with vaccine for up to 72 h, individual experiments shown. Reads are divided into ones with (blue) and without (red) mΨCmΨAG-sequence. P.values calculated using Wilcoxon test, p. value adjustment with Benjamini-Hochberg method.

**b,** Changes in poly(A) tails lengths for BMDMs WT (left panel) and *Tent5a*$^{flox/flox}$/*Tent5c*$^{-/-}$ (right panel) in time course following addition of mRNA-1273. Z-score normalized poly(A) lengths are represented in a colour scale.

**c,** Expression of genes induced in WT BMDMs upon mRNA-1273 addition, shown for WT (left panel) and *Tent5a*$^{flox/flox}$/*Tent5c*$^{-/-}$ cells (right panel). Expression values were z-score normalized.

**d,** Poly(A) lengths distribution of firefly luciferase spike-in (left panel) and mRNA-1273 (right panel), co-transfected to WT BMDMs. RNA was analyzed 24h post-transfection. Reads are divided into ones with (blue) and without (red) mΨCmΨAG sequence.

**e,** Western blots on spike protein and firefly luciferase co-transfected to WT BMDMs

**f,** Spike protein levels in serum of immunized mice, WT and *Tent5a*$^{-/-}$, measured 2 and 4 days after immunization. Statistics calculated with paired Wilcoxon test.

**g,** Total IgG levels in serum of immunized mice, WT and *Tent5a*$^{-/-}$, measured 14 days after immunization. Statistics calculated with Wilcoxon test.

### Extended Data Figure 6

**a**

#### Figure 2a

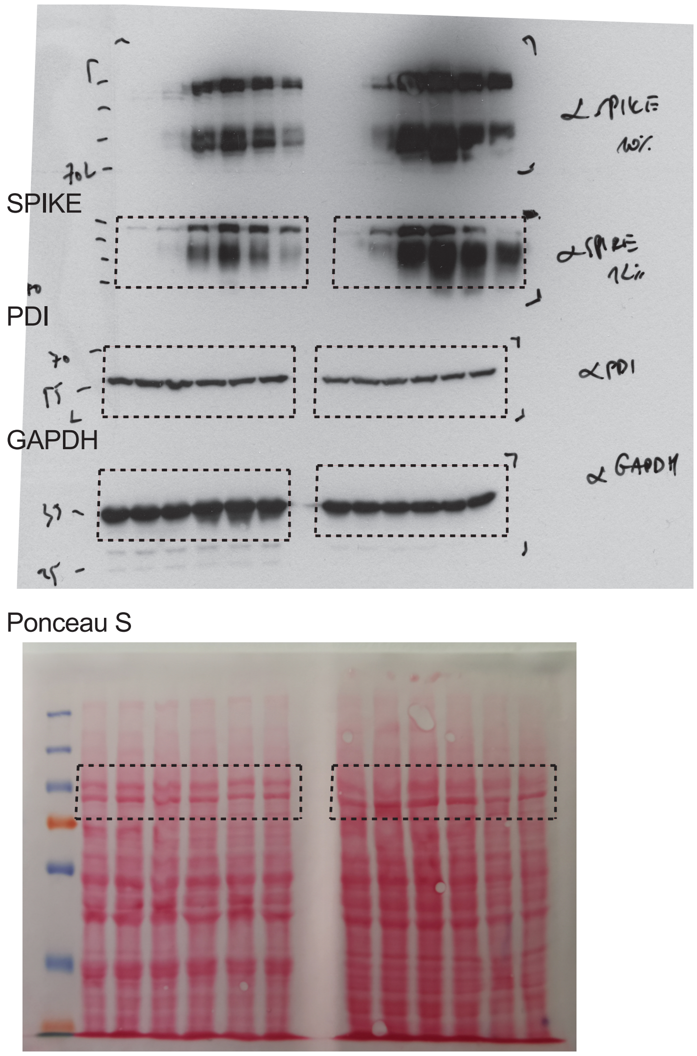


**b**

#### Figure 2d

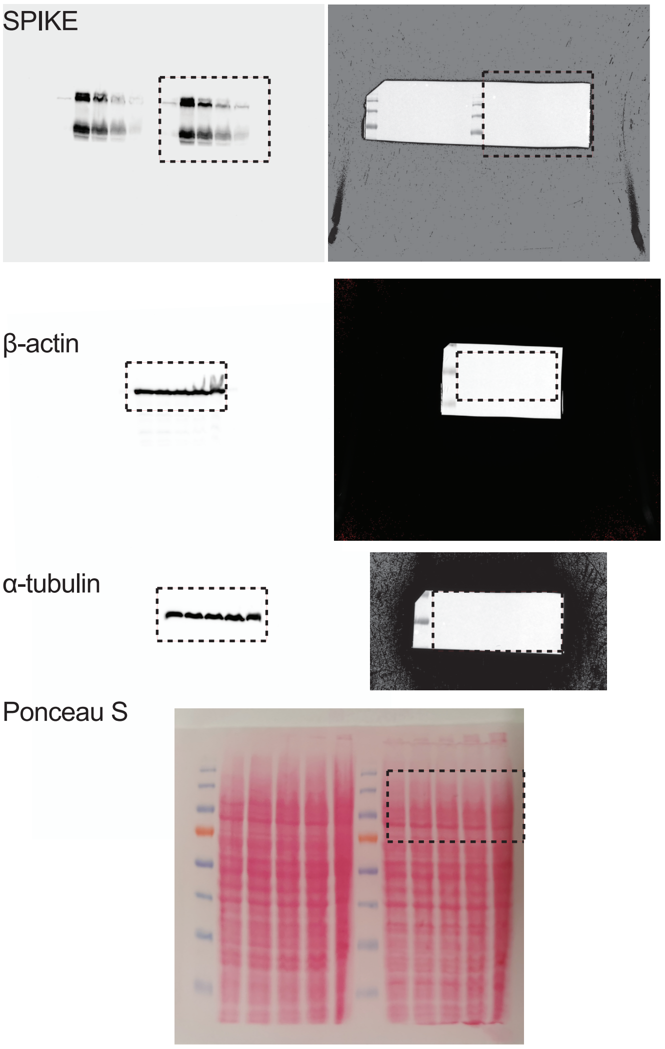


**c**

#### Figure 3d

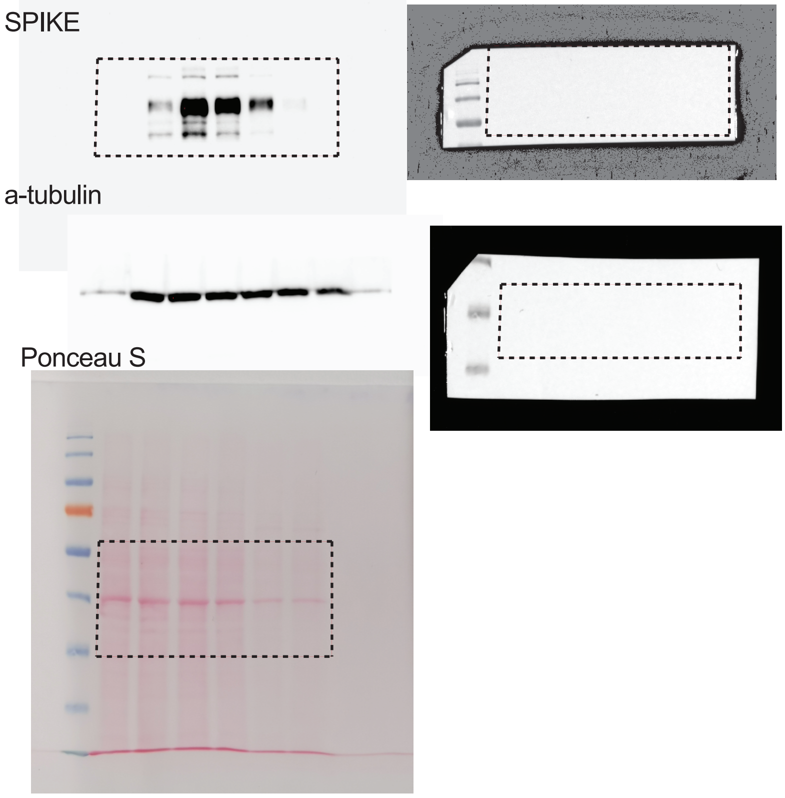


**d**

#### Extended Data Figure 5e

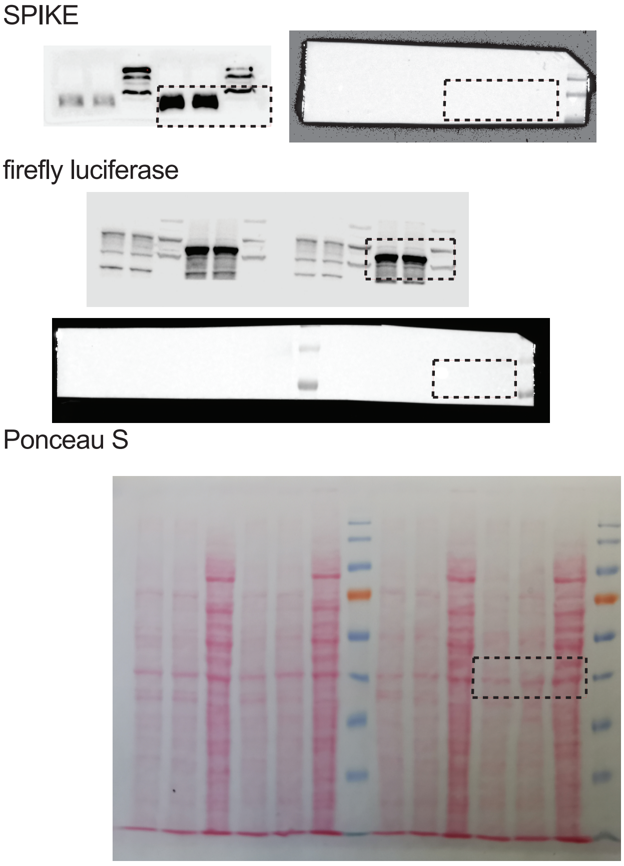

**Extended Data Figure 6. Uncropped blots (related to Figure 2a, 2d, 3d and Extended Data Figure 5e).**

**a,** Western blots and membrane Ponceau S staining corresponding to Fig. 2a.

**b,** Western blots and membrane Ponceau S staining corresponding to Fig. 2d.

**c,** Western blots and membrane Ponceau S staining corresponding to Fig. 3d.

**d,** Western blots and membrane Ponceau S staining corresponding to ED Fig. 5e.
