## Supplementary Tables legends for "SARS-CoV-2 mRNA vaccine is re-adenylated *in vivo*, enhancing antigen production and immune response"

**Supplementary Table 1** (separate file)

Differential expression of genes at mRNA-1273 injection sites after mice immunization. DESeq2 LRT test statistics and clustering of significantly changed genes shown.

**Supplementary Table 2** (separate file)

GO Terms (Biological Process) enriched for genes in clusters exhibiting clear patterns of expression change at mRNA-1273 injection sites after mice immunization.

**Supplementary Table 3** (separate file)

Differential expression of genes in WT BMDMs upon mRNA-1273 delivery. DESeq2 LRT test statistics and clustering of significantly changed genes shown.

**Supplementary Table 4** (separate file)

GO Terms (Biological Process) enriched for genes in clusters exhibiting clear patterns of expression change in WT BMDMs upon mRNA-1273 delivery.

**Supplementary Table 5** (separate file)

Differential expression of genes in DCs upon mRNA-1273 delivery. DESeq2 LRT test statistics shown.

**Supplementary Table 6** (separate file)

Mean poly(A) lengths for genes expressed in WT and $\textit{Tent5a}^{\textit{Flox/Flox}}/\textit{Tent5c}^{-/-}$ BMDMs. Statistic for change in poly(A) lengths was calculated using Kruskall-Wallis test for WT samples and genes with at least 20 reads in each group. Multiple testing correction (adjusted p. value) was done using Benjamini-Hochberg method. Efect size is calculated as rank epsilon squared. For transcripts with the most pronounced poly(A) change (adjusted p. value < 0.001) their cluster membership is indicated.

**Supplementary Table 7** (separate file)

GO Terms (Biological Process) enriched for genes in clusters exhibiting clear patterns of poly(A) length change in WT BMDMs upon mRNA-1273 delivery.

**Supplementary Table 8** (separate file)

Differential expression of genes in $\textit{Tent5a}^{\textit{Flox/Flox}}/\textit{Tent5c}^{-/-}$ BMDMs upon mRNA-1273 delivery. DESeq2 LRT test statistics shown, and WT BMDMs cluster membership is indicated.
